# GABA production induced by imipridones is a targetable and imageable metabolic alteration in diffuse midline gliomas

**DOI:** 10.1101/2024.06.07.597982

**Authors:** Georgios Batsios, Suresh Udutha, Céline Taglang, Anne Marie Gillespie, Benison Lau, Sunjong Ji, Timothy Phoenix, Sabine Mueller, Sriram Venneti, Carl Koschmann, Pavithra Viswanath

## Abstract

Diffuse midline gliomas (DMGs) are lethal primary brain tumors in children. The imipridones ONC201 and ONC206 induce mitochondrial dysfunction and have emerged as promising therapies for DMG patients. However, efficacy as monotherapy is limited, identifying a need for strategies that enhance response. Another hurdle is the lack of biomarkers that report on drug-target engagement at an early timepoint after treatment onset. Here, using ^1^H-magnetic resonance spectroscopy, which is a non-invasive method of quantifying metabolite pool sizes, we show that accumulation of γ-aminobutyric acid (GABA) is an early metabolic biomarker that can be detected within a week of ONC206 treatment, when anatomical alterations are absent, in mice bearing orthotopic xenografts. Mechanistically, imipridones activate the mitochondrial protease ClpP and upregulate the stress-responsive transcription factor ATF4. ATF4, in turn, upregulates glutamate decarboxylase, which synthesizes GABA, and downregulates *ABAT*, which degrades GABA, leading to GABA accumulation in DMG cells and tumors. Functionally, GABA secreted by imipridone-treated cells acts in an autocrine manner via the GABAB receptor to induce expression of superoxide dismutase (SOD1), which mitigates imipridone-induced oxidative stress and, thereby, curbs apoptosis. Importantly, blocking autocrine GABA signaling using the clinical stage GABAB receptor antagonist SGS-742 exacerbates oxidative stress and synergistically induces apoptosis in combination with imipridones in DMG cells and orthotopic tumor xenografts. Collectively, we identify GABA as a unique metabolic adaptation to imipridones that can be leveraged for non-invasive assessment of drug-target engagement and therapy. Clinical translation of our studies has the potential to enable precision metabolic therapy and imaging for DMG patients.

**One Sentence Summary:** Imipridones induce GABA accumulation in diffuse midline gliomas, an effect that can be leveraged for therapy and non-invasive imaging.

## INTRODUCTION

Diffuse midline gliomas (DMGs) are universally fatal primary brain tumors in children with no long-term survivors (*1, 2*). These tumors, which include those previously referred to as diffuse intrinsic pontine glioma, are molecularly driven by a mutation in histone H3.3 or H3.1 at lysine 27 (H3K27-altered DMGs) (*3, 4*). The H3K27M mutation results in global loss of trimethylation of H3K27 and reciprocal gain of H3K27 acetylation, leading to epigenetic changes that ultimately drive tumorigenesis (*3, 4*). DMGs arise in delicate anatomical areas such as the thalamus, brainstem, and spinal cord, which prevent total surgical resection (*1, 2*). Fractionated radiation is standard of care, but is largely palliative, and median overall survival remains 9-11 months after initial diagnosis (*5, 6*).

The imipridone ONC201 (dordaviprone) is a first-in-class, brain-penetrant, small molecule agent that has emerged as a promising therapeutic option for children with DMGs (*7, 8*). Originally identified as a dopamine receptor 2 (DRD2) antagonist (*9*), recent studies suggest that ONC201 also functions as an agonist of the ATP-dependent mitochondrial caseinolytic protease ClpP (*10–12*). ClpP forms a complex with ClpX that contributes to the maintenance of protein homeostasis by degrading misfolded proteins in the mitochondria (*12, 13*). ClpP agonism by imipridones causes degradation of mitochondrial proteins, including the electron transport chain components complexes I, II and IV, and activates the integrated stress response (*8, 10–14*). Mitochondria are the major source of reactive oxygen species (*15, 16*), and studies show that imipridones induce oxidative stress and apoptosis in cancer cells (*12–14, 17*). ONC201 is brain penetrant and significantly extends survival of mice bearing patient-derived DMG xenografts (*13, 17, 18*). ONC206 is a fluorinated derivative of ONC201 with the same core structure but ∼10-fold higher potency in growth inhibition assays against DMG cells (*19*). ONC206 also demonstrates efficacy in preclinical DMG models and is undergoing phase I clinical testing in DMG patients (*13*). Early pilot clinical studies showed that ONC201 has efficacy as monotherapy in adult and pediatric patients with H3K27M-mutant gliomas (*20–22*). Importantly, children with brainstem or thalamic H3K27M tumors treated with ONC201 prior to recurrence had a median overall survival of 21.7 months vs. 12 months for historical controls (*18*). Another study in patients with H3K27M DMGs treated with ONC201 after tumor recurrence following radiation reported a median overall survival of 13.7 months (*23*). While these findings of extended survival and clinical benefit conferred by ONC201 are undoubtedly significant (*24*), tumor regression following treatment remains elusive, highlighting the need for additional strategies that enhance response to imipridone therapy.

Metabolic reprogramming promotes tumor growth as well as resistance to therapy (*25*). Previous studies have delineated the metabolic effects of imipridone treatment on DMGs and other solid tumors (*13, 17, 18*). *In vitro* stable isotope tracing with ^13^C-labeled glucose or glutamine indicates that ONC201 suppresses glucose metabolism via the tricarboxylic acid (TCA) cycle while potentiating glutamine metabolism to α-ketoglutarate (α-KG) and L-2-hydroxyglutarate in DMG cells (*18*). L-2-hydroxyglutarate, which is a potent competitive inhibitor of α-KG-dependent histone demethylases, partially restores histone H3K27 trimethylation in tumor xenografts and in patient tissue (*18*). However, whether imipridone-induced alterations in glucose or glutamine metabolism can be leveraged to enhance response to therapy is unclear.

A significant hurdle in the development and clinical deployment of novel DMG therapies is the lack of pharmacodynamic biomarkers of drug-target engagement and response to therapy. Due to the deep-seated anatomical location within delicate midline areas of the brain, quantifying drug penetrability and drug-target engagement by examining serial biopsies is challenging. As a result, assessment of disease progression and response to therapy is heavily dependent on magnetic resonance imaging (MRI) (*26*). However, DMGs are diffusely infiltrative tumors with indistinct borders, which makes it difficult, even for experienced personnel, to accurately quantify tumor volume (*26*). Importantly, there is no consensus on what constitutes a biologically meaningful reduction in tumor volume following the onset of therapy and current response assessment in pediatric neuro-oncology guidelines require comparison of changes in serial MRI scans separated by at least 8 weeks (*26*).

This long delay has devastating consequences such as ineffective treatment and unnecessary toxicity for patients with progressive disease. It also hinders the proper interpretation of the efficacy of novel agents in clinical trials. Magnetic Resonance Spectroscopy (MRS) is an advanced MRI-based method used to quantify metabolism in cells, animals, and humans (*27, 28*). MRS generates spectra from tissue voxels in which each metabolite resonates at a distinct chemical shift that appears as a peak, and the peak integral corresponds to the concentration (*27, 28*). Due to the difference in tissue concentration between water (75 M) and metabolites (1-30 mM), the spatial resolution of MRS is lower than MRI. However, DMGs are often inoperable due to the anatomical location, and patients tend to present with substantial tumor burden. MRS, therefore, is ideal for interrogating the metabolic state of tumor tissue in DMGs *in vivo* in their natural microenvironment. ^1^H-MRS quantifies steady-state metabolite pool sizes (*27, 28*), which provide a readout of the balance between metabolite synthesis and turnover (*29, 30*). Importantly, ^1^H-MRS has been used in clinical research in DMG patients (*28*). We focused on delineating metabolic rewiring induced by imipridones in clinically relevant patient-derived and syngeneic DMG models, with the goal of identifying metabolic vulnerabilities that can be exploited for therapy and for non-invasive metabolic imaging. Our studies indicate that accumulation of the metabolic signaling molecule γ-aminobutyric acid (GABA) following imipridone treatment is an early ^1^H-MRS-detectable imaging biomarker of drug-target engagement in mice bearing orthotopic DMG xenografts at the clinically relevant magnetic field strength of 3T. Mechanistic studies show that imipridones upregulate GABA synthesis by glutamate decarboxylase (GAD1) and downregulate GABA catabolism by the GABA transaminase (ABAT), in a ClpP- and ATF4-dependent manner. Functionally, autocrine GABA signaling via the GABAB receptor drives expression of superoxide dismutase 1 (SOD1), which scavenges superoxide radicals and curbs apoptosis induced by imipridones. Importantly, genetic or pharmacological inhibition of the GABAB receptor synergistically induces cell death in combination with ONC206, thereby identifying a novel therapeutic opportunity.

## RESULTS

### Imipridones induce a ^1^H-MRS-detectable increase in GABA pool size in patient-derived and syngeneic DMG cells

To determine whether imipridones induce imageable metabolic alterations, we interrogated the effect of treatment with ONC201 or ONC206 on ^1^H-MRS-detectable metabolite pool sizes in DMG cells. To this end, we examined patient-derived H3.3 K27M (BT245, SF8628), syngeneic H3.3 K27M (26-B7) or H3.1 K27M (26-C2) DMG cells. First, consistent with prior studies (*13*), we confirmed that ONC201 and ONC206 inhibited the viability of DMG cells with IC50 values of ∼1.5 μM for ONC201 and ∼0.2 μM for ONC206 (Fig. 1A-1B). We then treated DMG cells with vehicle or ONC206 and examined the effect on steady-state metabolite levels using ^1^H-MRS. As shown in the representative ^1^H-MRS spectra in Fig. 1C and Fig. S1A-S1C and the quantification in Fig. 1D, GABA was uniquely detectable in ONC206-treated cells but not vehicle-treated controls in all DMG models. Concomitantly, glutamate was significantly elevated while succinate and aspartate were significantly reduced in ONC206-treated cells relative to vehicle-treated controls in all DMG models (Fig. 1E-1H). We observed a significant reduction in citrate, methionine, glutathione, phosphocholine, glycine, and myoinositol in some but not all models (Fig. 1E-1H). Of note, we confirmed that treatment with ONC201 also increases ^1^H-MRS-detectable GABA in our DMG models (Fig. 1I). To orthogonally confirm these results and to determine whether GABA is secreted out of the cell, we used an enzyme-linked immunosorbent assay (ELISA) to measure intracellular and secreted GABA concentrations in imipridone-treated cells. As shown in Fig. 1J-1K, ONC206 and ONC201 significantly upregulated both intracellular and secreted GABA pool sizes in DMG cells. Collectively, these results point to GABA accumulation as a unique metabolic consequence of imipridone treatment that can be detected by ^1^H-MRS in patient-derived and syngeneic DMG cells.

**Fig. 1.**
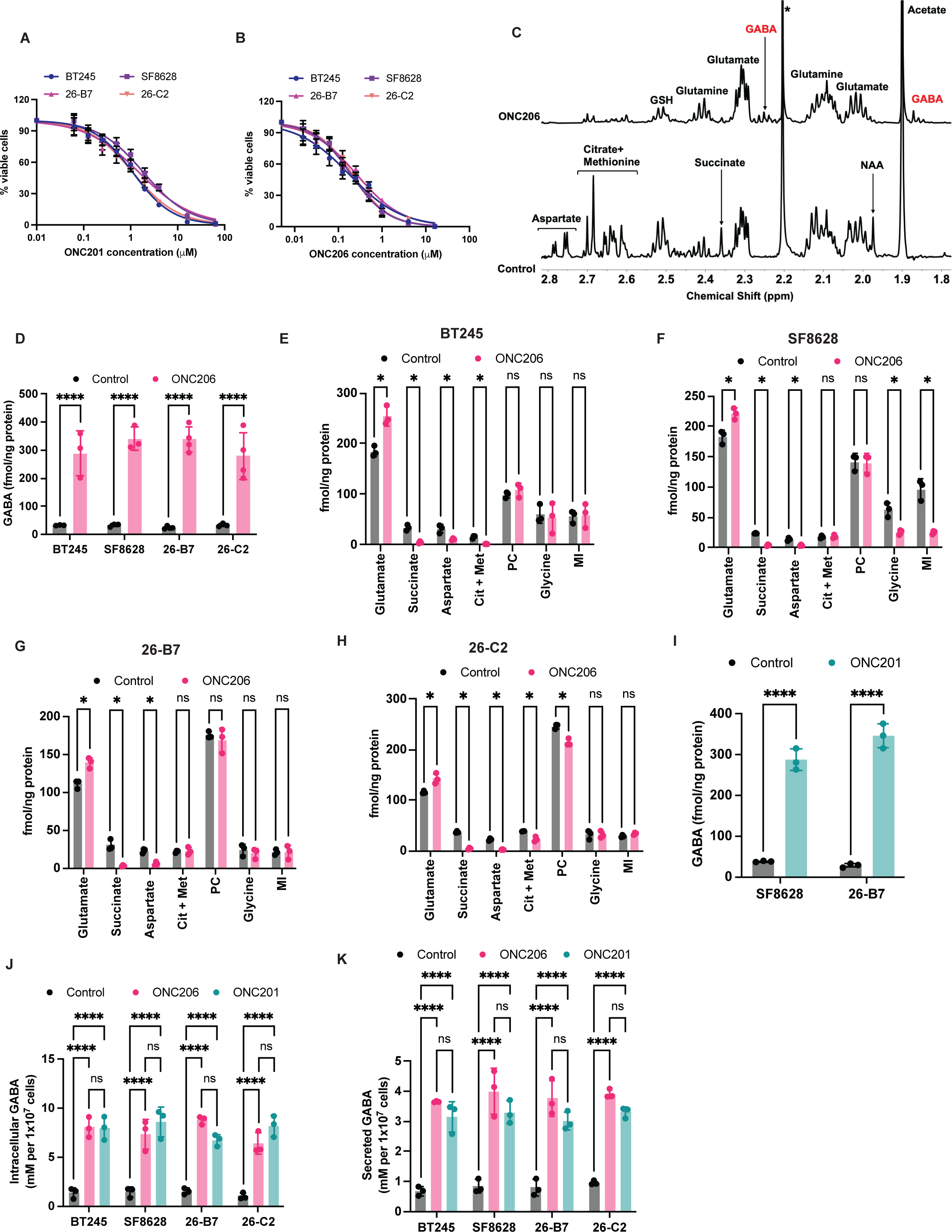
^1^H-MRS identifies GABA as a unique metabolic biomarker in patient-derived and syngeneic DMG cells. Dose response curves for ONC201 **(A)** and ONC206 **(B)** in BT245, SF8628, 26-B7 and 26-C2 cells. Cells were treated with the indicated concentrations of the drugs for 72 h and % live cells calculated using the CellTiter-Fluor™ Cell Viability Assay. **(C)** Representative ^1^H-MRS spectra from BT245 cells treated with vehicle (DMSO) or 500 nM ONC206 for 72 h. * indicates a peak for acetone. **(D)** Quantification of ^1^H-MRS-detectable GABA pool size in BT245, SF8628, 26-B7 and 26-C2 cells treated with vehicle (DMSO) or 500 nM ONC206 for 72 h. Quantification of ^1^H-MRS-detectable glutamate, succinate, aspartate, citrate + methionine (Cit + Met, composite peak), phosphocholine (PC), glycine and myoinositol (MI) in BT245 **(E)**, SF8628 **(F)**, 26-B7 **(G)** and 26-C2 **(H)** cells treated with vehicle (DMSO) or 500 nM ONC206 for 72 h. **(I)** Quantification of ^1^H-MRS-detectable GABA pool size in SF8628, and 26-B7 cells treated with vehicle (DMSO) or 10 μM ONC201 for 72 h. Intracellular **(J)** and secreted **(K)** GABA concentration measured by ELISA in BT245, SF8628, 26-B7 and 26-C2 cells treated with vehicle (DMSO), 500 nM ONC206, or 10 μM ONC201 for 72 h.

### 1H-MRS-detectable GABA is elevated in mice bearing orthotopic patient-derived or syngeneic DMG xenografts at an early timepoint following ONC206 treatment

Obtaining an early readout of drug-target engagement following the onset of therapy is a significant problem in DMG patient management (*26*). Since our *in vitro* studies above identified a significant increase in GABA in imipridone-treated cells, we questioned whether an early increase in GABA served as a non-invasive imaging biomarker of drug-target engagement in DMG-bearing mice *in vivo* at clinical magnetic field strength (3T). To this end, we treated mice bearing intracranial patient-derived (BT245 or SF8628) xenografts with vehicle or ONC206 for a week and assessed tumor volume by MRI or optical imaging (bioluminescence) and tumor metabolism by ^1^H-MRS at 3T. Mice were then euthanized, and tissue resected for *ex vivo* assays of drug-target engagement (see experimental schematic in Fig. 2A). Previous studies indicate that ONC201 and ONC206 function as agonists of the mitochondrial protease ClpP, which forms a complex with ClpX (*10–12*). ClpP agonism causes degradation of ClpX and key proteins including the mitochondrial electron transport chain complex I, leading to activation of the integrated stress response, and the onset of apoptosis *in vitro* and *in vivo* (*10–13*). As shown in Fig. 2B-2C, expression of ClpX and complex I was lost while the integrated stress response transcription factor ATF4 was upregulated in ONC206-treated tumor tissue relative to vehicle-treated controls for both the SF8628 and BT245 models. Concomitantly, caspase activity was significantly elevated in ONC206-treated tumors relative to controls (Fig. 2D). Importantly, using the orthogonal ELISA, we confirmed that GABA concentration was significantly higher in ONC206-treated tumor tissue relative to vehicle-treated controls (Fig. 2E). These results provide molecular confirmation of drug-target engagement within a week of ONC206 treatment in mice bearing intracranial DMG xenografts *in vivo*.

**Fig. 2.**
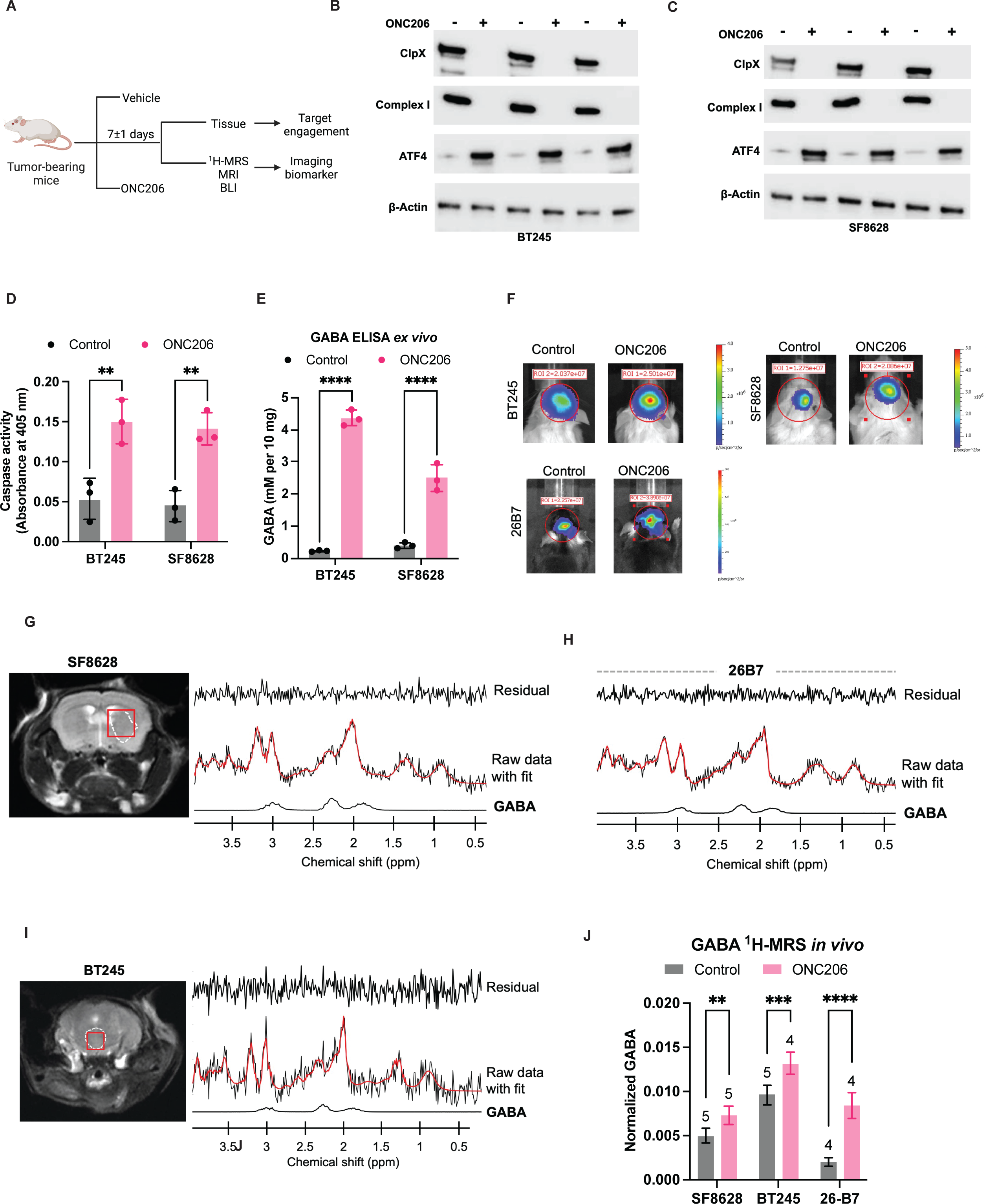
^1^H-MRS-detectable GABA is an early imaging biomarker of drug-target engagement in mice bearing orthotopic DMG xenografts *in vivo*. (A) Schematic representation of the experimental plan for *in vivo* and *ex vivo* studies. Western blots for ClpX, complex I, and ATF4 in tumor tissue resected from mice bearing orthotopic BT245 **(B)** or SF8628 **(C)** xenografts. β-actin was used as loading control. Tumor-bearing mice were treated with vehicle (saline) or ONC206 (25 mg/kg, twice daily). On day 7±1, mice were euthanized, and tumor tissue was resected for analysis. Caspase activity **(D)** and GABA concentration measured by ELISA **(E)** in tumor tissue from mice bearing orthotopic BT245 or SF8628 tumor xenografts treated with vehicle or ONC206 as described above. **(F)** Representative bioluminescent images from mice bearing orthotopic BT245, SF8628 or 26-B6 xenografts treated with vehicle or ONC206 as described above. Data was acquired on day 7±1 on an IVIS scanner. **(G)** Representative ^1^H-MRS data acquired from a mouse bearing an orthotopic SF8628 tumor at day 7±1 after treatment with ONC206 (25 mg/kg, twice daily). The T2-weighted MRI with the tumor contoured in white dotted lines is shown on the left. The right panel shows the ^1^H-MRS spectrum with the fit from LCModel (red), the fit for GABA and the residual trace. **(H)** Representative ^1^H-MRS spectrum with the fit from LCModel (red), the fit for GABA and the residual trace acquired from a mouse bearing an orthotopic 26-B7 tumor at day 7±1 after treatment with ONC206 (25 mg/kg, twice daily). **(I)** Representative ^1^H-MRS data acquired from a mouse bearing an orthotopic BT245 tumor implanted in the pons at day 7±1 after treatment with ONC206 (25 mg/kg, twice daily). The T2-weighted MRI with the tumor contoured in white dotted lines is shown on the left. The right panel shows the ^1^H-MRS spectrum with the fit from LCModel (red), the fit for GABA and the residual trace. **(J)** Quantification of *in vivo* ^1^H-MRS-detectable GABA from mice bearing orthotopic BT245, SF8628, or 26-B7 tumors treated with vehicle (saline) or ONC206 (25 mg/kg, twice daily). On day 7±1, ^1^H-MRS data was acquired on a Bruker 3T scanner. For the bar graph, the numbers above each bar indicate the number of mice in each group.

Examination of tumor volume by T2-weighted MRI (Fig. S2A) or optical imaging (Fig. 2F) did not identify differences at day 7±1 between DMG-bearing mice that were treated with vehicle or ONC206 in any of our models. We then performed *in vivo* single-voxel (3×3×3 mm^3^) ^1^H-MRS using a PRESS sequence at 3T on mice bearing intracranial patient-derived (BT245, SF8628) and syngeneic (26-B7) tumor xenografts at this early timepoint (day 7±1) after treatment with ONC206. As shown in the representative spectra in Fig. 2G-2I, GABA was clearly detectable by ^1^H-MRS at day 7±1 in all DMG models *in vivo.* Importantly, quantification of the data (Fig. 2J) confirmed a significant increase in GABA at this early timepoint (day 7±1) in ONC206-treated mice relative to vehicle-treated controls for all models. We did not observe consistent changes in ^1^H-MRS-detectable levels of myo-inositol + glycine, glutamate + glutamine (glx), or total choline *in vivo* across our models (Fig. S2B-S2D). In line with prior studies, ONC206 significantly (albeit modestly) improved survival of mice bearing intracranial SF8628 tumors (49 days for vehicle-treated controls vs. 62 days for ONC206-treated mice, p<0.05; Fig. S2E). Collectively, these findings indicate that elevated ^1^H-MRS-detectable GABA provides a non-invasive readout of drug-target engagement following imipridone treatment that can be visualized at an early timepoint when anatomical changes are absent in mice bearing intracranial DMG xenografts *in vivo*.

### Imipridones upregulate glutamate decarboxylase (GAD1) and downregulate GABA transaminase (ABAT) in a ClpP- and ATF4-dependent manner in DMGs

To identify the metabolic source of elevated GABA, we examined the effect of imipridones on pathways of GABA synthesis and catabolism in DMG cells. As shown in Fig. 3A, GABA is generated via decarboxylation of glutamate. Glutamate can be directly generated from glutamine or from α-KG produced by oxidation of glucose-derived pyruvate via the TCA cycle. GABA is, in turn, catabolized to succinate semialdehyde and then succinate, which enters the TCA cycle to generate ATP and NADH. To determine whether glucose or glutamine is the source of elevated GABA, we performed liquid chromatography mass spectrometry (LC-MS) following stable isotope tracing with [U-^13^C]-glucose or [U-^13^C]-glutamine in BT245, SF8628, 26-B7, and 26-C2 cells treated with vehicle or ONC206. As shown in Fig. 3B-3E, we observed a significant increase in ^13^C labeling of glutamate, α-KG, 2-hydroxyglutarate, and GABA from [U-^13^C]-glutamine in ONC206-treated DMG cells. Concomitantly, we observed a significant decrease in ^13^C labeling of succinate and aspartate from [U-^13^C]-glutamine in all DMG models (Fig. 3B-3E). We did not observe alterations in [U-^13^C]-glucose metabolism to glutamate, GABA, or succinate, while ^13^C labeling of α-KG, fumarate, malate, aspartate, and citrate from [U-^13^C]-glucose was reduced following ONC206 treatment in DMG cells (Fig. S2F-S2I). We confirmed that the pool size of GABA, was elevated in ONC206-treated cells in all DMG models (Fig. 3F), further confirming our ^1^H-MRS and ELISA results. Taken together, these findings identify glutamine-derived glutamate as the precursor of GABA in imipridone-treated DMG cells.

**Fig. 3.**
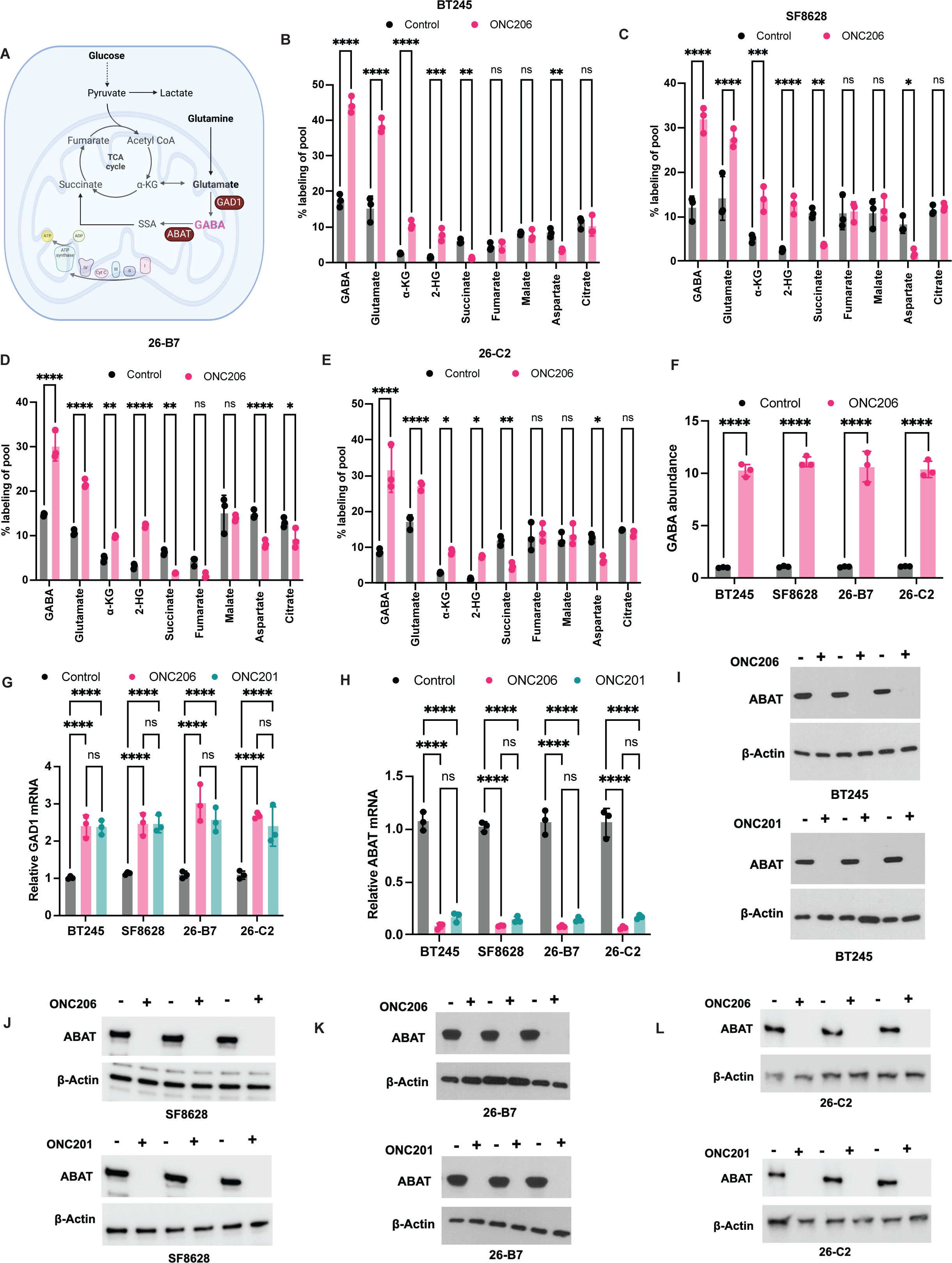
Glutamine is the metabolic source of GABA in imipridone-treated DMG cells. >**(A)** Schematic representation of pathways of GABA synthesis from glucose or glutamine. Quantification of % ^13^C labeling of metabolites from [U-^13^C]-glutamine in BT245 **(B)**, SF8628 **(C)**, 26-B7 **(D)** and 26-C2 **(E)** cells treated with vehicle (DMSO) or 500 nM ONC206 for 72 h. **(F)** GABA abundance (pool size) measured by LC-MS in BT245, SF8628, 26-B7 and 26-C2 cells treated with vehicle (DMSO) or 500 nM ONC206 for 72 h. Expression of GAD1 **(G)** and ABAT **(H)** measured by QPCR in BT245, SF8628, 26-B7 and 26-C2 cells treated with vehicle (DMSO), 500 nM ONC206, or 10 μM ONC201 for 72 h. Western blots for ABAT in in BT245 **(I)**, SF8628 **(J)**, 26-B7 **(K)** and 26-C2 **(L)** cells treated with vehicle (DMSO), 500 nM ONC206, or 10 μM ONC201 for 72 h. In each case, the top panel indicates blots from cells treated with vehicle or ONC206 and the bottom panel indicates blots from cells treated with vehicle or ONC201. β-actin was used as loading control.

Since the pool size of a metabolite depends on the balance between synthesis and degradation, we examined the effect of imipridones on the expression of GAD1, the enzyme which catalyzes the conversion of glutamate to GABA, and ABAT, which converts GABA to succinate semialdehyde (see schematic in Fig. 3A). As shown in Fig. 3G-3L, while ONC201 and ONC206 upregulated expression of GAD1, they completely abrogated expression of ABAT at both mRNA and protein levels in all DMG models (BT245, SF8628, 26-B7, and 26-C2).

We then focused on the molecular mechanism by which imipridones modulate expression of GAD1 and ABAT in DMG cells. As demonstrated by previous studies (*7, 10, 13*) and our data above (see Fig. 2B-2C), ClpP agonism by imipridones activates expression of the stress-responsive transcription factor ATF4. Since ATF4 is known to regulate expression of genes involved in amino acid metabolism (*31, 32*), we hypothesized that ATF4 upregulation by ClpP modulates expression of GAD1 and ABAT in imipridone-treated DMGs. To test this hypothesis, we examined the effect of silencing ClpP (Fig. S3A) or ATF4 (Fig. S3B) using pooled CRISPR sgRNA on GAD1 and ABAT expression in DMG cells. As shown in Fig. 4A-4F and Fig. S3C-S3F, silencing ClpP or ATF4 reduced expression of GAD1 and restored expression of ABAT in imipridone-treated BT245, SF8628, 26-B7 and 26-C2 cells. Conversely, overexpressing ATF4 (see Fig. S3G for confirmation of overexpression) upregulated GAD1 and downregulated ABAT in vehicle-treated DMG cells (Fig. 4G-4L). Importantly, chromatin immunoprecipitation quantitative PCR (ChIP-QPCR) confirmed that ATF4 binds to the GAD1 and ABAT promoters in imipridone-treated cells (Fig. 4M-4N), further confirming the mechanistic link between ATF4 and expression of GAD1 and ABAT. Collectively, these results suggest that activation of ATF4 modulates expression of GAD1 and ABAT in imipridone-treated DMG cells.

**Fig. 4.**
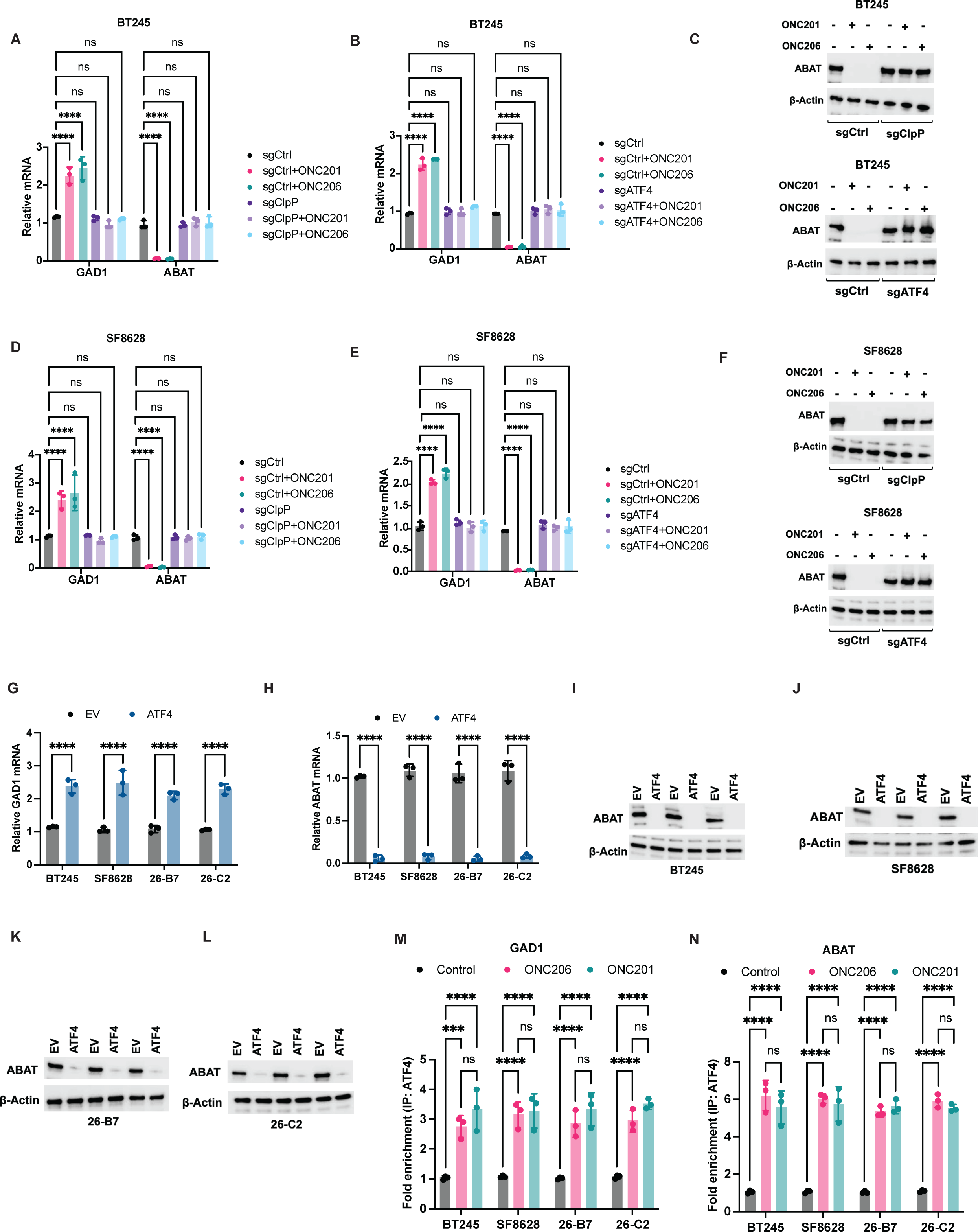
Imipridones upregulate GAD1 and downregulate ABAT in a ClpP- and ATF4-dependent manner in DMG cells. Expression of GAD1 and ABAT measured by QPCR in BT245 cells expressing sgRNA against ClpP **(A)** or ATF4 **(B)**. Cells expressing a scrambled sgRNA sequence were used as control (sgControl). Cells were treated with vehicle (DMSO), 500 nM ONC206, or 10 μM ONC201 for 72 h. **(C)** Western blots for ABAT in BT245 sgControl, sgClpP (top panel) or sgATF4 (bottom panel) cells treated with vehicle (DMSO), 500 nM ONC206, or 10 μM ONC201 for 72 h. Expression of GAD1 and ABAT measured by QPCR in SF8628 cells expressing sgRNA against ClpP **(D)** or ATF4 **(E)**. Cells expressing a scrambled sgRNA sequence were used as control (sgControl). Cells were treated with vehicle (DMSO), 500 nM ONC206, or 10 μM ONC201 for 72 h. **(F)** Western blots for ABAT in SF8628 sgControl, sgClpP (top panel) or sgATF4 (bottom panel) cells treated with vehicle (DMSO), 500 nM ONC206, or 10 μM ONC201 for 72 h. Expression of GAD1 **(G)** and ABAT **(H)** measured by QPCR in BT245, SF8628, 26-B7 and 26-C2 cells transfected with an empty vector (EV) or a plasmid expressing ATF4 (ATF4). Western blots for ABAT in BT245 **(I)**, SF8628 **(J)**, 26-B7 **(K)** and 26-C2 **(L)** cells transfected with an empty vector or a plasmid expressing ATF4. Quantification of ATF4 binding to the GAD1 **(M)** and ABAT **(N)** promoters measured by ChIP-QPCR in BT245, SF8628, 26-B7 and 26-C2 cells treated with vehicle (DMSO), 500 nM ONC206, or 10 μM ONC201 for 72 h.

### Imipridones upregulate GAD1 and downregulate ABAT in DMG tumor xenografts

To confirm our findings above *in vivo*, we examined tumor tissue resected at day 7±1 from mice bearing orthotopic SF8628 or BT245 tumor xenografts that were treated with vehicle or ONC206 as described in Fig. 2A. GAD1 expression was significantly elevated (Fig. 5A-5C) while ABAT expression was significantly downregulated (Fig. 5D-5F) in ONC206-treated tumor tissue relative to vehicle-treated tumors in both the SF8628 and BT245 models. Concomitantly, ATF4 binding to the GAD1, and ABAT promoters was significantly higher in ONC206-treated tumor tissue relative to vehicle-treated tumors (Fig. 5G-5H). These results are consistent with the increase in GABA observed by *in vivo* ^1^H-MRS (Fig. 2J) and by ELISA (Fig. 2E) in ONC206-treated tumor tissue relative to controls. They are also consistent with the loss of ClpX and upregulation of ATF4 in ONC206-treated tumor tissue (see Fig. 2B-2C). Taken together, our findings suggest that imipridones drive GABA accumulation by upregulating GAD1 and downregulating ABAT in DMG cells and orthotopic tumor xenografts.

**Fig. 5.**
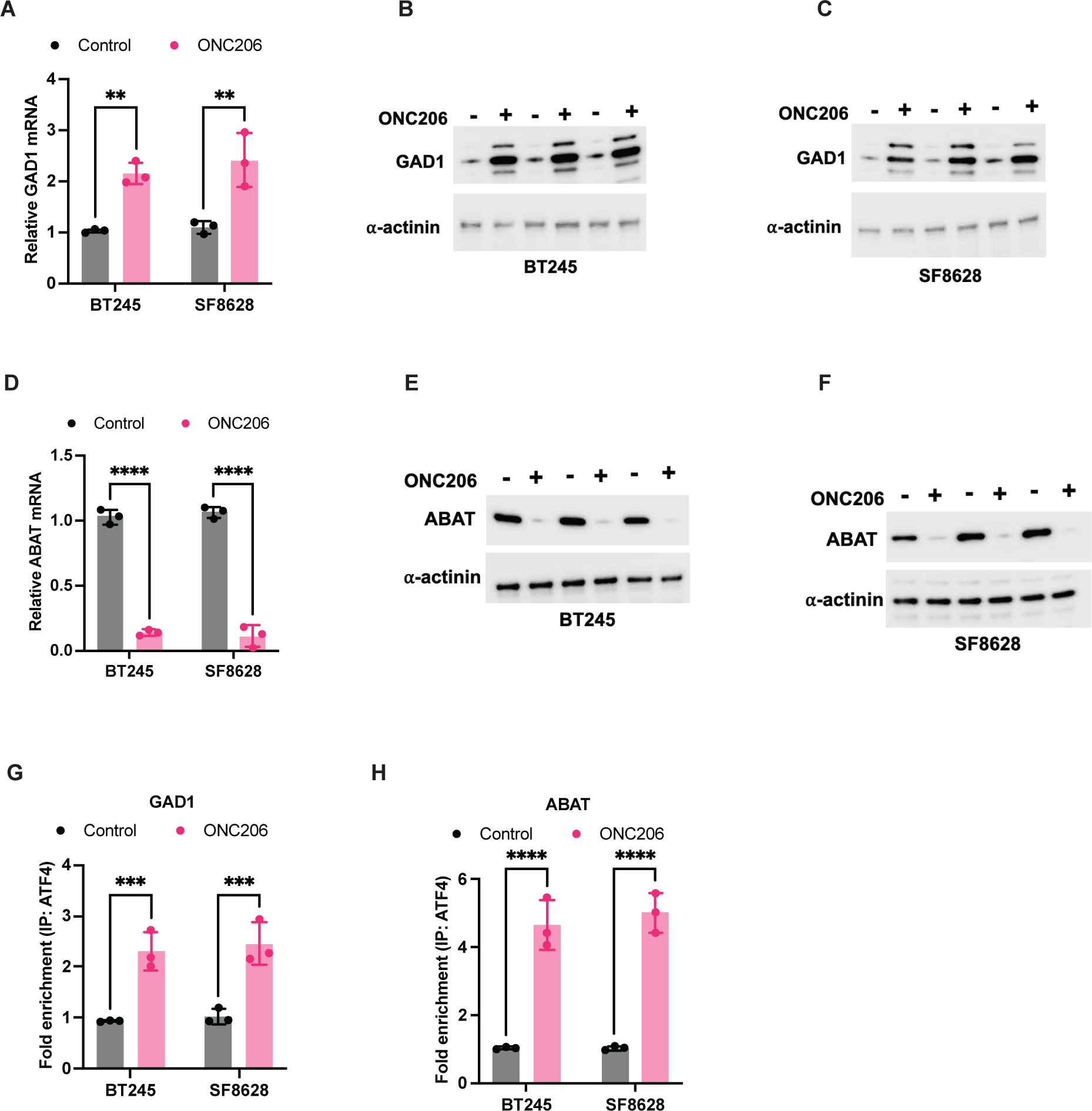
Imipridones upregulate GAD1 and downregulate ABAT in DMG tumors *in vivo*. >**(A)** GAD1 mRNA measured by QPCR in tumor tissue resected from mice bearing orthotopic BT245 or SF8628 tumor xenografts treated with vehicle (saline) or ONC206 (25 mg/kg, twice daily). On day 7±1, mice were euthanized, and tumor tissue was resected for analysis. GAD1 protein measured by western blotting in tumor tissue resected from mice bearing orthotopic BT245 **(B)** or SF8628 **(C)** tumor xenografts treated with vehicle or ONC206 as described above. **(D)** ABAT mRNA measured by QPCR in tumor tissue resected from mice bearing orthotopic BT245 or SF8628 tumor xenografts treated with vehicle (saline) or ONC206 (25 mg/kg, twice daily). On day 7±1, mice were euthanized, and tumor tissue was resected for analysis. ABAT protein measured by western blotting in tumor tissue resected from mice bearing orthotopic BT245 **(E)** or SF8628 **(F)** tumor xenografts treated with vehicle or ONC206 as described above. Quantification of ATF4 binding to the GAD1 **(G)** and ABAT **(H)** promoters measured by ChIP-QPCR in tumor tissue resected from mice bearing orthotopic BT245 or SF8628 tumor xenografts treated with vehicle or ONC206 as described above.

### Imipridone-induced GABA accumulation is a metabolic adaptation that mitigates oxidative stress and curbs apoptosis in DMG cells

To assess the functional consequences of imipridone-induced GABA accumulation in DMGs, we first examined the effect of overexpressing ABAT or an empty vector (see Fig. S4A-S4D for validation of overexpression) on DMG cells treated with vehicle or imipridones. We confirmed that ABAT overexpression depleted both intracellular and secreted GABA in BT245, SF8628, 26-B7 or 26-C2 cells treated with ONC201 or ONC206, returning it to levels observed in vehicle-treated DMG cells (Fig. 6A-6D and Fig. S4E-S4H). Previous studies have linked GABA to oxidative stress signaling in mammalian and plant cells (*33–36*). Mitochondria are the major source of reactive oxygen species and defects in electron transport chain components are associated with oxidative stress (*15, 16, 37*). Since ClpP agonism by imipridones degrades mitochondrial proteins, including complex I (see Fig. 2I-2J) and induces oxidative stress (*10, 12–14, 17*), we examined the effect of depleting GABA by overexpressing ABAT on imipridone-induced oxidative stress in DMG cells. As shown in Fig. 6E-6H and Fig. S4I-S4L, imipridones caused accumulation of superoxide radicals, and induced apoptosis in DMG cells, consistent with prior studies (*10, 12–14, 17*). Importantly, depleting GABA potentiated superoxide radical generation, and apoptosis in DMG cells (Fig. 6E-6H, Fig. S4I-S4L). To further confirm these results, we examined the effect of the ABAT inhibitor vigabatrin on DMG cells. We confirmed that vigabatrin increased pool sizes of intracellular and secreted GABA in DMG cells (Fig. S5A-S5B). Importantly, as shown in Fig. 6I-6L and Fig. S5C-S5F, vigabatrin alleviated oxidative stress and blocked apoptosis induced by imipridones in DMG cells.

**Fig. 6.**
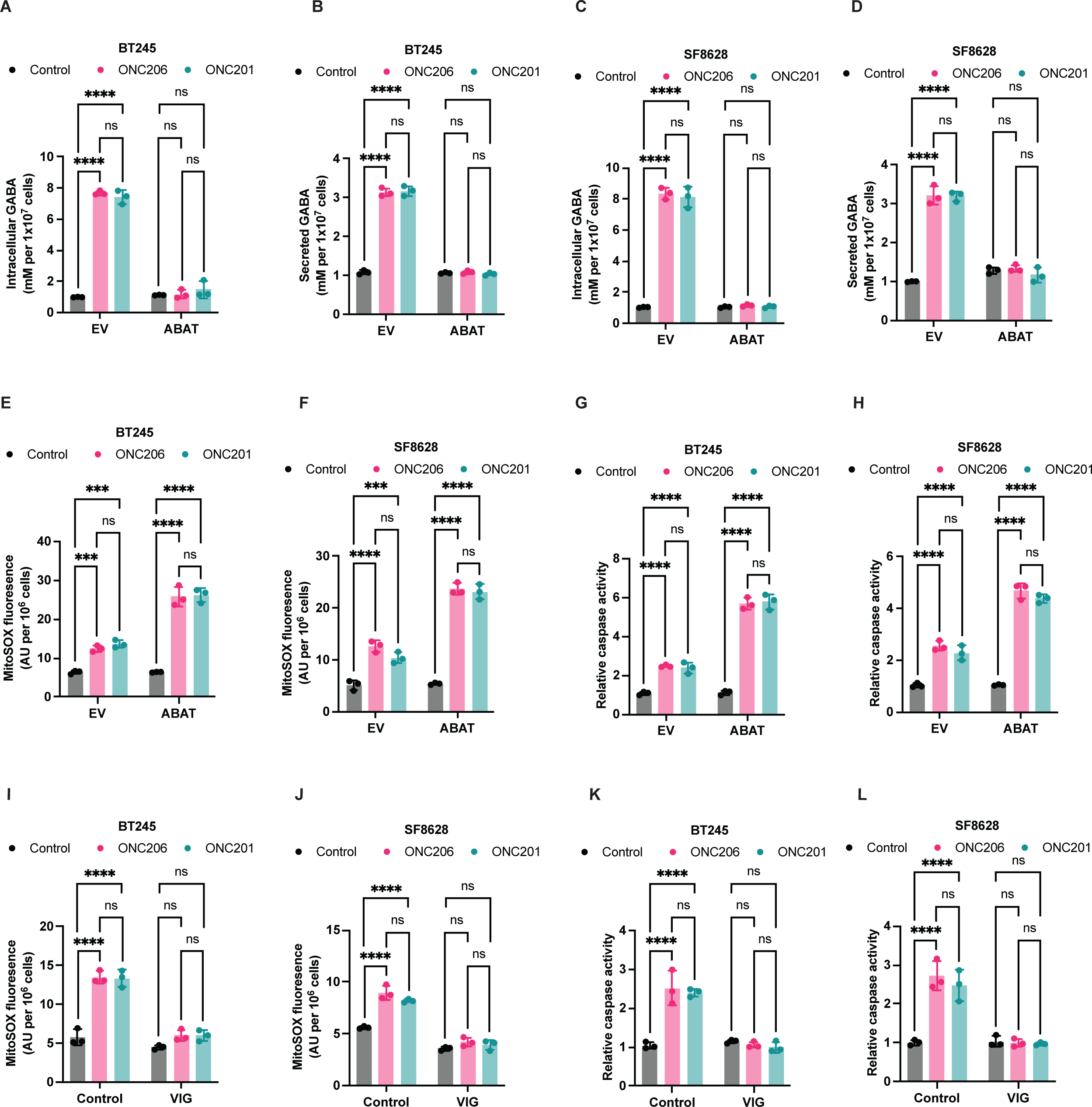
GABA mitigates oxidative stress and curbs apoptosis induced by imipridones in DMG cells. Intracellular **(A)** and secreted **(B)** GABA concentration in BT245 cells transfected with an empty vector (EV) or a plasmid expressing ABAT (ABAT). Cells were treated with vehicle (DMSO), 500 nM ONC206, or 10 μM ONC201 for 72 h. Intracellular **(C)** and secreted **(D)** GABA concentration in SF8628 cells transfected with an empty vector (EV) or a plasmid expressing ABAT (ABAT). Cells were treated with vehicle (DMSO), 500 nM ONC206, or 10 μM ONC201 for 72 h. Superoxide radical levels in BT245 **(E)** and SF8628 **(F)** cells transfected with an empty vector (EV) or a plasmid expressing ABAT (ABAT). Cells were treated with vehicle (DMSO), 500 nM ONC206, or 10 μM ONC201 for 72 h. Caspase activity measured using the Caspase-Glo® 3/7 3D Assay in BT245 **(G)** and SF8628 **(H)** cells transfected with an empty vector (EV) or a plasmid expressing ABAT (ABAT). Cells were treated with vehicle (DMSO), 500 nM ONC206, or 10 μM ONC201 for 72 h. Superoxide radical levels in BT245 **(I)** and SF8628 **(J)** cells treated with vehicle (DMSO) or vigabatrin (VIG; 100 μM). Cells were concurrently treated with vehicle (DMSO), 500 nM ONC206, or 10 μM ONC201 for 72 h. Caspase activity measured using the Caspase-Glo® 3/7 3D Assay in BT245 **(K)** and SF8628 **(L)** cells treated with vehicle (DMSO) or vigabatrin (VIG; 100 μM). Cells were concurrently treated with vehicle (DMSO), 500 nM ONC206, or 10 μM ONC201 for 72 h.

Superoxide dismutase (SOD1) is the key enzyme that regulates oxidative stress by converting superoxide radicals to oxygen and hydrogen peroxide, which is subsequently detoxified to oxygen and water (*38*). We confirmed that imipridones upregulated SOD1 expression at both mRNA and protein levels in DMG cells, an effect that was normalized by ABAT overexpression (Fig. 7A-7H). In contrast, the ABAT inhibitor vigabatrin enhanced SOD1 expression in imipridone-treated DMG cells (Fig. 7I-7L). Collectively, these results suggest that GABA accumulation induced by imipridones alleviates oxidative stress and curbs apoptosis in DMG cells by driving SOD1 expression.

**Fig. 7.**
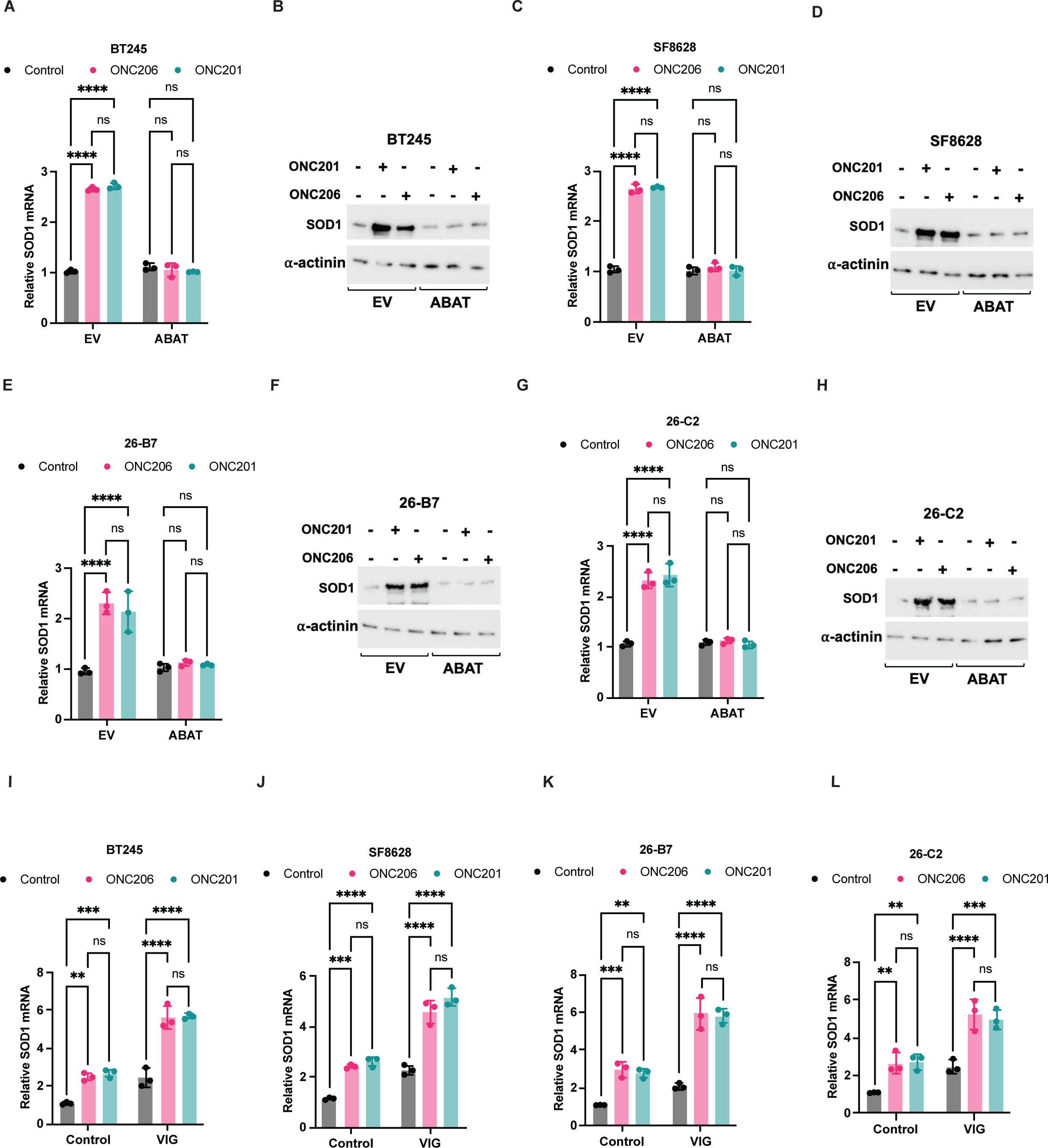
GABA upregulates SOD1 in imipridone-treated DMG cells. SOD1 mRNA **(A)** and protein **(B)** in BT245 cells transfected with an empty vector (EV) or a plasmid expressing ABAT (ABAT). Cells were treated with vehicle (DMSO), 500 nM ONC206, or 10 μM ONC201 for 72 h. SOD1 mRNA **(C)** and protein **(D)** in SF8628 cells transfected with an empty vector (EV) or a plasmid expressing ABAT (ABAT). Cells were treated with vehicle (DMSO), 500 nM ONC206, or 10 μM ONC201 for 72 h. SOD1 mRNA **(E)** and protein **(F)** in 26-B7 cells transfected with an empty vector (EV) or a plasmid expressing ABAT (ABAT). Cells were treated with vehicle (DMSO), 500 nM ONC206, or 10 μM ONC201 for 72 h. SOD1 mRNA **(G)** and protein **(H)** in 26-C2 cells transfected with an empty vector (EV) or a plasmid expressing ABAT (ABAT). Cells were treated with vehicle (DMSO), 500 nM ONC206, or 10 μM ONC201 for 72 h. SOD1 mRNA in BT245 **(I),** SF8628 **(J)**, 26-B7 **(K)** and 26-C2 **(L)** cells treated with vehicle (DMSO) or vigabatrin (VIG; 100 μM). Cells were concurrently treated with vehicle (DMSO), 500 nM ONC206, or 10 μM ONC201 for 72 h.

### GABA activates autocrine signaling via the GABAB receptor in DMG cells

Next, we examined the mechanism by which GABA activates SOD1 expression in DMG cells. The biological effects of GABA are largely mediated via signaling cascades triggered by GABA receptors (*39, 40*), and recent studies have highlighted a role for autocrine GABA signaling via these receptors in promoting tumor growth (*39–44*). Since our results above suggested that GABA was secreted out of the cell, we investigated whether autocrine GABA receptor signaling regulated SOD1 expression in DMG cells. There are two distinct classes of GABA receptors: GABAA and GABAB (*40*). The GABAA receptor is a ligand-gated chloride anion channel composed of five subunits while GABAB is a G-protein coupled receptor composed of a functional ligand-binding GABBR1 subunit and a structural GABBR2 subunit (*40*). Recent studies indicate that genes coding for GABAA receptor subunits are highly expressed in DMG cells (*45*). Here, by analyzing data from the Pediatric cBioportal, we confirmed that GABBR1 and GABBR2 are both highly expressed in H3K27M-altered tumors relative to high-grade gliomas that are H3 wild-type (Fig. 8A-8B). To assess if there is a causal relationship between imipridones and GABA receptors, we examined the effect of treatment with the GABAA receptor antagonist bicuculline or the clinical stage GABAB receptor antagonist SGS-742 on DMG cells (*46, 47*). Blocking GABAA receptor signaling using bicuculline did not normalize SOD1 expression in imipridone-treated cells in DMG cells (Fig. S6A-S6B). Importantly, as shown in Fig. 8C-8H and Fig. S6C-S6D, the GABAB receptor antagonist SGS-742 phenocopied GABA depletion and abrogated the increase in SOD1 expression caused by imipridones in the SF8628, BT245, 26-B7 and 26-C2 models. Furthermore, genetic ablation of GABBR1 using pooled sgRNA (see Fig. S6E for confirmation of GABBR1 silencing) also normalized SOD1 expression in imipridone-treated DMG cells (Fig. S6F-S6H).

**Fig. 8.**
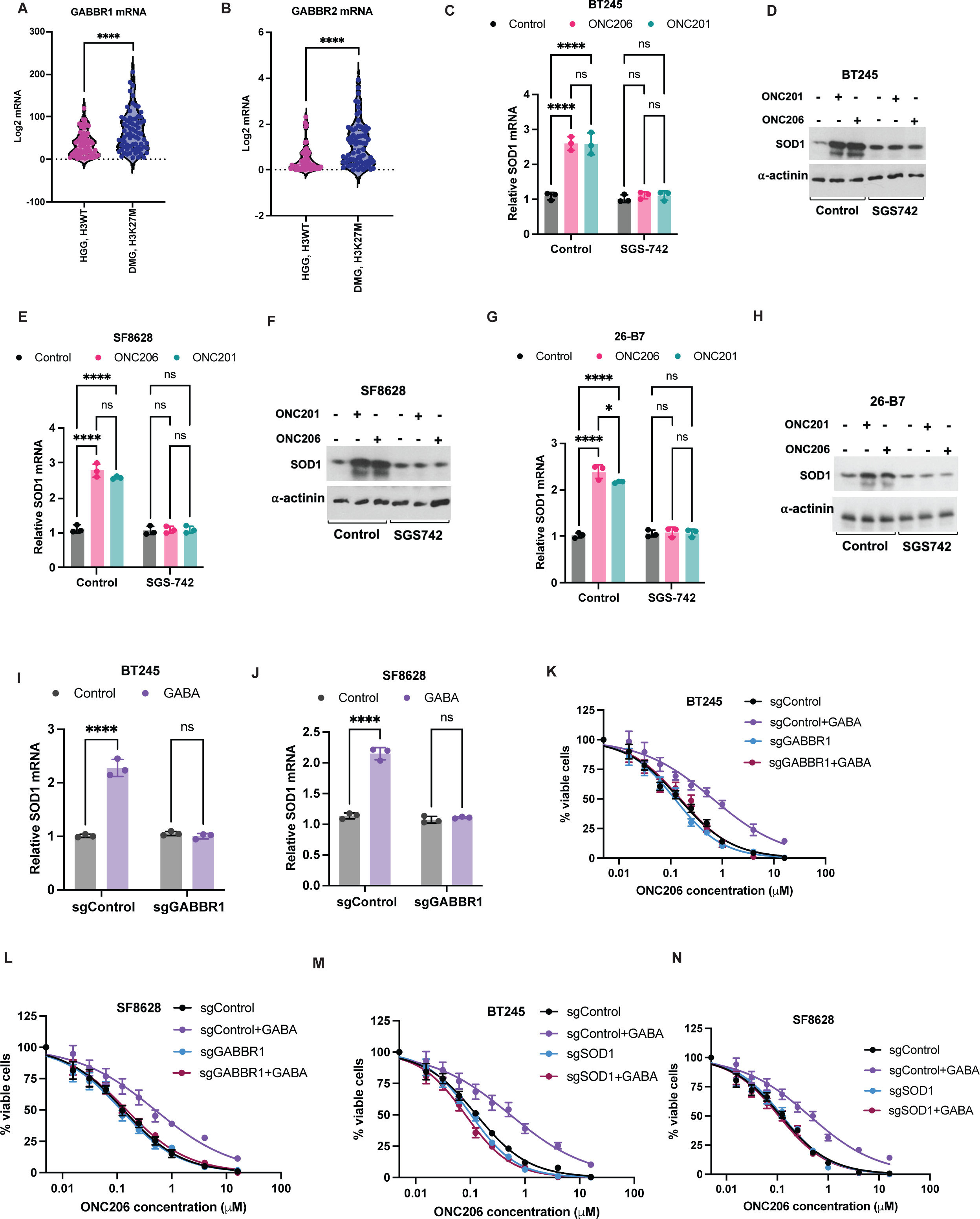
GABA activates SOD1 expression by autocrine signaling via the GABAB receptor in imipridone-treated DMG cells. mRNA levels of GABBR1 **(A)** and GABBR2 **(B)** in biopsies from DMG patients with H3K27-altered tumors or H3 wild-type tumors. Data was downloaded from the Open Pediatric Cancer Atlas at the Pediatric cBioportal. SOD1 mRNA **(C)** and protein **(D)** in BT245 cells treated with vehicle (DMSO) or SGS-742 (50 μM). Cells were concurrently treated with vehicle (DMSO), 500 nM ONC206, or 10 μM ONC201 for 72 h. SOD1 mRNA **(E)** and protein **(F)** in SF8628 cells treated with vehicle (DMSO) or SGS-742 (50 μM). Cells were concurrently treated with vehicle (DMSO), 500 nM ONC206, or 10 μM ONC201 for 72 h. SOD1 mRNA **(G)** and protein **(H)** in 26-B7 cells treated with vehicle (DMSO) or SGS-742 (50 μM). Cells were concurrently treated with vehicle (DMSO), 500 nM ONC206, or 10 μM ONC201 for 72 h. SOD1 mRNA in BT245 **(I)** and SF8628 **(J)** cells expressing scrambled (sgControl) or GABBR1-targeted sgRNA (sgGABBR1). Dose response curves for ONC206 in BT245 **(K)** and SF8628 **(L)** cells expressing scrambled (sgControl) or GABBR1-targeted sgRNA (sgGABBR1). Cells were concurrently treated with vehicle (saline) or 1 mM GABA for 72 h. Dose response curves for ONC206 in BT245 **(M)** and SF8628 **(N)** cells expressing scrambled (sgControl) or SOD1-targeted sgRNA (sgSOD1). Cells were concurrently treated with vehicle (saline) or 1 mM GABA for 72 h.

To mechanistically link GABA to GABAB receptor signaling and SOD1, we examined the effect of treatment with exogenous GABA on DMG cells with (sgControl) or without GABBR1 (sgGABBR1) expression. As shown in Fig. 8I-8J, GABA induced SOD1 expression in sgControl BT245 and SF8628 cells. Concomitantly, as shown in Fig. 8K-8L, GABA increased the IC50 of ONC206 by ∼4-5-fold in sgControl BT245 cells and SF8628 cells. Importantly, blocking autocrine GABA signaling by silencing GABBR1 normalized SOD1 expression and restored sensitivity to ONC206 in the presence of GABA for both BT245 and SF8628 models (Fig. 8I-8L). We also confirmed that silencing SOD1 phenocopied GABBR1 silencing and restored sensitivity to ONC206 in the presence of GABA for both BT245 and models (Fig. 8M-8N). Taken together, these studies indicate that autocrine GABA signaling via the GABAB receptor upregulates SOD1 and reduces the sensitivity of DMG cells to imipridones.

### Targeting the GABAB receptor synergistically induces cell death in combination with imipridones in DMG cells and orthotopic tumor xenografts

Given the mechanistic association between GABA signaling, SOD1, and imipridone-induced apoptosis, we questioned whether targeting the GABAB receptor would synergize with imipridone treatment. As shown in Fig. 9A-9D, using a high-throughput assay for caspase activity, we confirmed that blocking GABAB receptor signaling using SGS-742 synergized with ONC206 in inducing apoptosis of DMG cells (Bliss synergy scores of 17.2 for BT245 and 16.8 for SF8628; scores >10 indicate synergy). Concomitantly, superoxide radical generation was significantly higher in BT245, SF8628, 26-B7 and 26-C2 cells treated with the combination of SGS-742 and ONC201/ONC206 relative to ONC201/ONC206 as monotherapy (Fig. 9E-9H). We confirmed that genetic ablation of GABBR1 also potentiated superoxide radical production and apoptosis induced by ONC201 or ONC206 in DMG cells (Fig. 9I-9L).

**Fig. 9.**
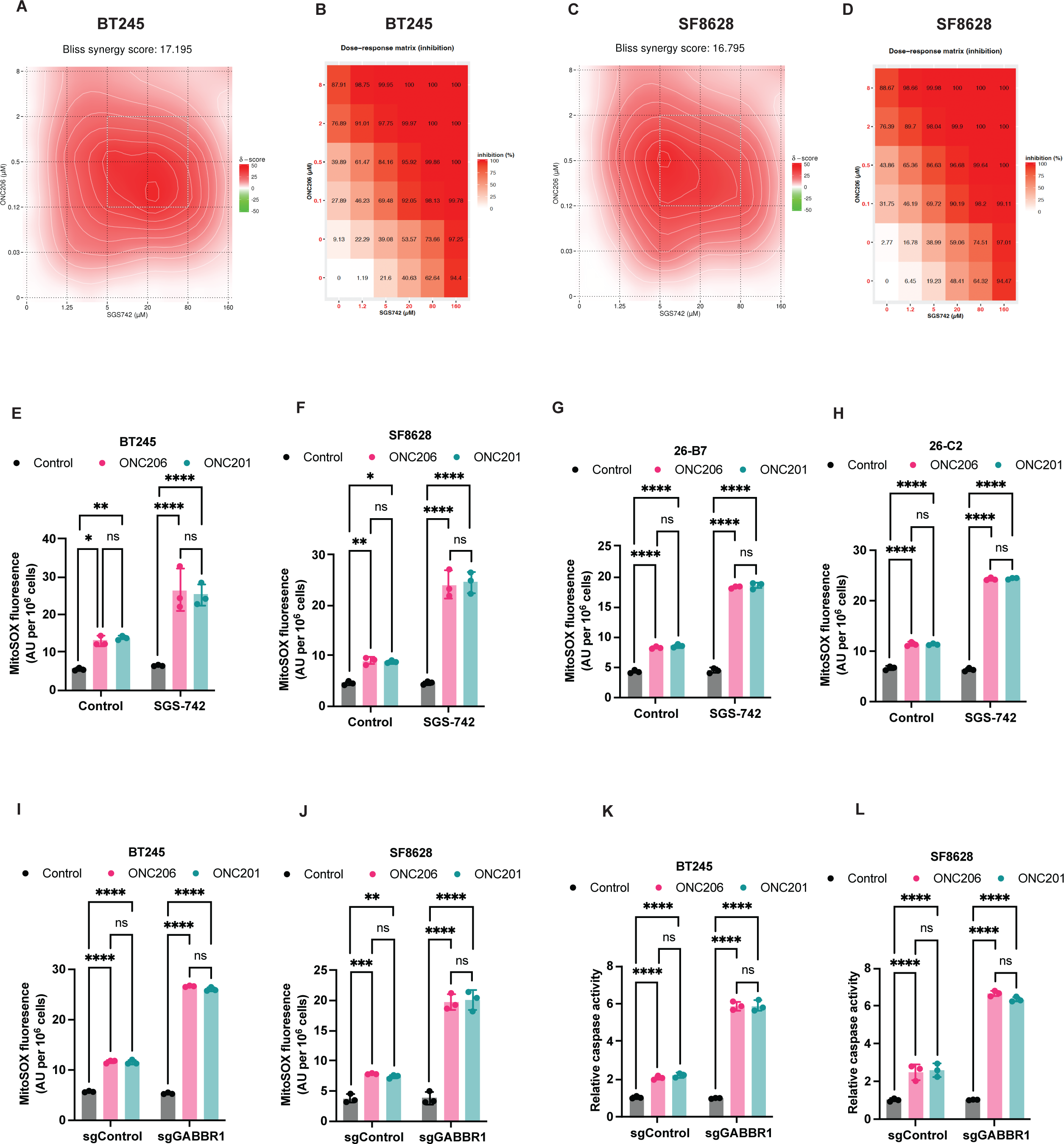
SGS-742 synergistically induces apoptosis in combination with imipridones in DMG cells. Bliss synergy map **(A)** and dose-response inhibition matrix **(B)** for the combination of SGS-742 and ONC206 in BT245 cells. Cells were treated with the indicated concentrations of the drugs for 72 h and % apoptotic cells calculated using the Caspase-Glo® 3/7 3D Assay. Bliss synergy map **(C)** and dose-response inhibition matrix **(D)** for the combination of SGS-742 and ONC206 in SF8628 cells. Cells were treated with the indicated concentrations of the drugs for 72 h and % apoptotic cells calculated using the Caspase-Glo® 3/7 3D Assay. Superoxide radical levels measured in BT245 **(E)**, SF8628 **(F)**, 26-B7 **(G)** and 26-C2 **(H)** cells treated with vehicle (DMSO) or SGS-742 (50 μM). Cells were concurrently treated with vehicle (DMSO), 500 nM ONC206, or 10 μM ONC201 for 72 h. Superoxide radical levels measured in BT245 **(I)** and SF8628 **(J)** cells expressing scrambled (sgControl) or GABBR1-targeted sgRNA (sgGABBR1). Cells were concurrently treated with vehicle (DMSO), 500 nM ONC206, or 10 μM ONC201 for 72 h. Caspase activity measured using the Caspase-Glo® 3/7 3D Assay in BT245 **(K)** and SF8628 **(L)** cells expressing scrambled (sgControl) or GABBR1-targeted sgRNA (sgGABBR1). Cells were concurrently treated with vehicle (DMSO), 500 nM ONC206, or 10 μM ONC201 for 72 h.

Next, we examined the therapeutic potential of inhibiting the GABAB receptor using SGS-742 in combination with imipridones in DMGs *in vivo.* To this end, we treated mice bearing intracranial SF8628 tumor xenografts with ONC206, alone or combined with SGS-742, for 7±1 days and resected tumor tissue for *ex vivo* assays (see experimental schematic in Fig. 10A). As shown in Fig. 10B-10D, while SOD1 expression and activity were elevated in tumors treated with ONC206 alone, combining SGS-742 with ONC206 normalized SOD1 expression and activity, returning it to levels observed in vehicle-treated tumors. Importantly, consistent with our *in vitro* data, caspase activity was significantly higher in tumors treated with the combination of SGS-742 and ONC206 relative to ONC206 alone (Fig. 10E). Collectively, these findings provide proof-of-principle evidence that targeting autocrine GABAB receptor signaling potentiates imipridone-induced cell death in mice bearing orthotopic DMG xenografts *in vivo*.

**Fig. 10.**
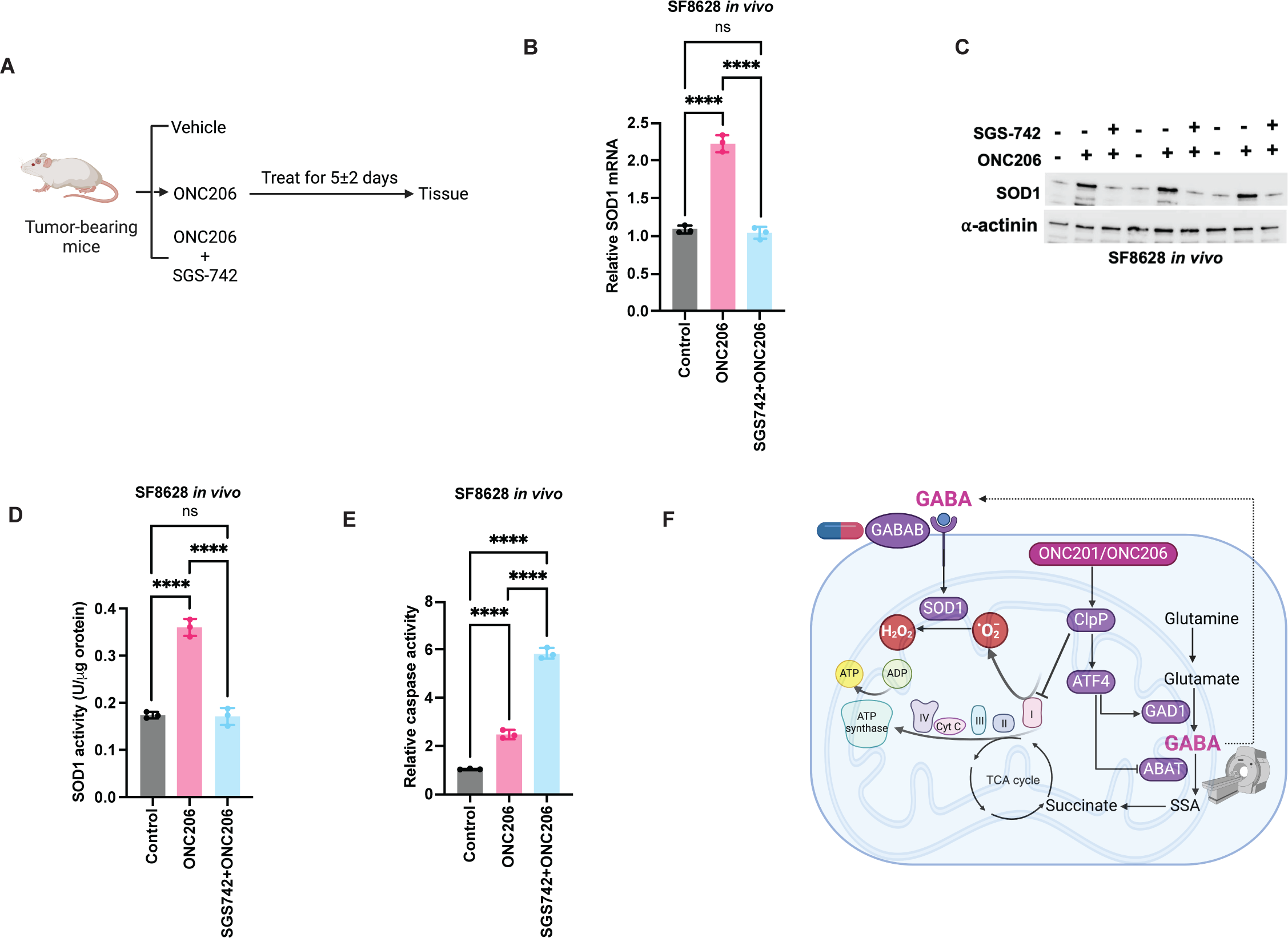
SGS-742 synergistically induces apoptosis in combination with imipridones in mice bearing orthotopic DMG xenografts. >**(A)** Schematic representation of the experimental plan for *in vivo* studies. SOD1 mRNA **(B)**, SOD1 protein **(C)**, SOD1 activity **(D)** and caspase activity **(E)** in tumor tissue resected from mice bearing orthotopic SF8628 tumor xenografts treated with vehicle (saline), ONC206 (25 mg/kg, twice daily), or the combination of ONC206 (25 mg/kg, twice daily) and SGS-742 (8 mg/kg, twice daily). On day 7±1, mice were euthanized, and tumor tissue was resected for analysis. **(F)** Schematic representation of the mechanistic consequences of imipridone-induced GABA accumulation in DMGs. ClpP agonism by ONC201 or ONC206 activates ATF4, which then upregulates GAD1 and downregulates ABAT, leading to accumulation of GABA. ^1^H-MRS enables visualization of this increase in GABA non-invasively at an early timepoint when changes are not detectable by anatomical imaging. GABA secreted out of the cell acts in an autocrine manner via the GABAB receptor to upregulate expression of SOD1. SOD1 scavenges superoxide radicals induced by imipridone treatment and partially rescues apoptosis in DMG cells. Blocking GABA signaling using the clinical stage GABAB receptor antagonist SGS-742 synergistically induces cell death in combination with imipridones in DMG cells and orthotopic tumor xenografts.

## DISCUSSION

DMGs are highly malignant brain tumors in children (*1, 2*). ONC201 confers meaningful clinical benefit with a significant extension in median overall survival in DMG patients (*18, 23, 24*). ONC206 is a structural analog of ONC201 with higher potency in preclinical DMG models and is currently in clinical trials (*7, 13, 19*). While these results are significant and serve to highlight the possibility of developing meaningful therapies for DMGs, nevertheless, patients succumb to the disease within ∼27 months. Therefore, there is a need to understand the molecular effects of imipridones and, in particular, identify adaptive strategies that limit the efficacy of these promising drugs. Here, using multiple approaches i.e., ^1^H-MRS, LC-MS and ELISA, we show that imipridones induce GABA accumulation in patient-derived and syngeneic murine DMG models. Importantly, imipridone-induced GABA can be non-invasively visualized *in vivo* using ^1^H-MRS. GABA acts in an autocrine manner via the GABAB receptor to upregulate SOD1, which then mitigates oxidative stress and reduces apoptosis induced by imipridones in DMG cells and orthotopic tumor xenografts. Targeting autocrine GABA signaling using the GABAB receptor inhibitor SGS-742 synergistically induces cell death in combination with imipridones in DMG cells. Importantly, SGS-742 normalizes SOD1 expression and potentiates ONC206-induced apoptosis *in vivo*. Collectively, as summarized in Fig. 10E, we identify GABA as an imipridone-induced metabolic adaptation, potential therapeutic target, and non-invasive imaging biomarker in preclinical DMG models *in vitro* and *in vivo*.

Our studies highlight, for the first time, the utility of ^1^H-MRS for metabolic imaging of drug-target engagement following imipridone treatment in DMG cells and orthotopic tumor xenografts. In this context, it is important to note that the increase in GABA provides positive contrast at an early timepoint after imipridone treatment when changes cannot be observed by anatomical imaging. With regard to clinical translation, ^1^H-MRS has been used in clinical research in DMG patients (*28*). Notably, ^1^H-MRS is the only biomedical imaging modality capable of visualizing GABA *in vivo* (*48–51*). ^1^H-MRS has been used to detect GABA in the human brain, including in children, under healthy and pathological conditions (*48–51*). Of particular relevance is the fact that our studies were performed using a standard PRESS sequence at the widely available clinical magnetic field strength of 3T. The ease of clinical translation of our findings is likely to be further enhanced by the increased availability of specialized acquisition sequences such as GABA-edited ^1^H-MRS and denoising techniques that improve the signal to noise ratio and reduce scan times (*48–51*). Given the increasing use of imipridones for treatment of DMG patients, our studies provide a strong rationale for clinical integration of ^1^H-MRS into routine longitudinal assessment of DMG patients treated with imipridones.

We identify a role for the stress-responsive transcription factor ATF4 in driving GABA accumulation in imipridone-treated DMG cells and tumors. Although originally identified as dopamine receptor DRD2 antagonists, studies suggest that imipridones are allosteric agonists of the ClpP protease, which is highly expressed in DMGs (*9–13*). We show that ClpP agonism by imipridones upregulates ATF4 in DMG cells and tumor xenografts. ATF4, in turn, upregulates GAD1 and downregulates ABAT, thereby driving GABA accumulation in DMG cells and tumors. Silencing ClpP or ATF4 normalizes expression of GAD1 and ABAT and depletes GABA in imipridone-treated DMG cells. In light of the observation that inactivating mutations in ClpP confer resistance to ONC201 (*12, 18*), our findings raise the intriguing possibility that ^1^H-MRS-based quantification of GABA may report on the emergence of resistance to imipridones *in vivo*.

GABA is best known for its role as the principal inhibitory neurotransmitter in the central nervous system (*52, 53*). GABA synthesized and released from GABAergic neurons exerts its effects on neuronal excitability and function by signaling via the GABAA or GABAB receptors, and dysregulation of GABAergic signaling is implicated in neurological disorders such as epilepsy and stroke (*52, 53*). In this context, it is important to note that recent cancer neuroscience studies suggest that GABAergic neurons form synapses with DMG cells and signal via the GABAA receptors to promote DMG growth in preclinical models (*45*). Our studies indicate that imipridone-induced GABA acts in an autocrine manner via the GABAB receptor to alleviate oxidative stress in tumor cells. Further studies are needed to assess whether GABA also influences neurotransmission or glioma-neuronal interactions. In a similar vein, studies have also identified a role for tumor-derived GABA in promoting immunosuppression by excluding T cells from the tumor microenvironment in preclinical cancer models (*44*). GABA can also suppress anti-tumor immunity by inducing interleukin-10 production that reprograms tumor-associated macrophages to an anti-inflammatory state (*54*). We anticipate that future studies will focus on determining whether GABA secreted into the tumor microenvironment following imipridone treatment promotes immunosuppression, and whether this can be therapeutically leveraged to enhance anti-tumor immunity.

To the best of our knowledge, our studies are the first to identify a role for GABA in driving SOD1 expression in cancer. By activating ClpP, imipridones induce degradation of components of the mitochondrial electron transport chain, including complex I, II and IV (*7, 10–14*). Defects in complex I are involved in the increased production of reactive oxygen species, including superoxide radicals, in neurodegenerative diseases (*15, 16, 37*). Indeed, imipridones induce production of superoxide radicals in cancer cells, including DMGs (*12–14, 17*). Beyond its role as a neurotransmitter, GABA is best known for its role in alleviating oxidative stress in plant and mammalian cells (*33–36*). Studies show that GABA protects human umbilical vein endothelial cells against damage induced by reactive oxygen species (*35*). GABA has also been shown to protect pancreatic β cells against oxidative stress and apoptosis (*36*). These effects of GABA were mediated via modulation of glycogen synthase kinase-3β or NRF2 signaling (*35, 36*). GABA signaling via GABAA or GABAB receptors interfaces with downstream pathways, including glycogen synthase kinase-3β, the MAPK/ERK, PI3K/Akt, or β-catenin signaling, all of which play key roles in regulating tumor proliferation, and apoptosis (*33, 39, 40*). Our studies point to minimal GABA production in untreated DMG cells and indicate that imipridones cause accumulation of GABA which is secreted out of the cell. Using genetic and pharmacological tools for depleting or potentiating GABA levels, we show that GABA acts via the GABAB receptor to induce SOD1 expression and mitigate oxidative stress in imipridone-treated cells. We further confirmed that treatment with exogenous GABA reduces DMG sensitivity to ONC206, an effect that is rescued by GABAB receptor inhibition. Although further studies are needed to identify the signaling pathway linking GABAB receptor activation to SOD1, nonetheless, our studies identify an imipridone-GABA-SOD1 signaling network in DMGs. The net effect of this GABAergic signaling cascade is partial rescue of apoptosis induced by imipridones, thereby, hindering the efficacy of these drugs.

Our studies highlight the therapeutic potential of targeting GABA signaling to enhance response to imipridones in DMGs. We show that inhibiting GABAB using SGS-742 results in strong synergy with ONC206 *in vitro.* Importantly, combination therapy with SGS-742 and ONC206 significantly potentiates ONC206-induced apoptosis in mice bearing orthotopic tumor xenografts *in vivo*. From the standpoint of clinical translation, SGS-742 is a brain-penetrant, selective GABAB receptor antagonist (*47*). It does not inhibit GABAA or other receptors in the central nervous system (*47*). Preclinical studies in mice, rats, and monkeys and a phase 2 study in adults with mild cognitive impairment showed that oral administration of SGS-742 was well-tolerated and significantly improved attention, reaction time, visual information processing, and working memory (*47, 55, 56*). Of note, SGS-742 was recently tested in children with succinic semialdehyde deficiency, which leads to abnormal accumulation of the downstream GABA metabolite γ-hydroxybutyric acid. Although SGS-742 failed to improve cognition in this setting, it should be noted that there were no drug-related toxicities in children (*57*). These prior studies, taken together with our data, provide a rationale for clinical testing of the combination of SGS-742 and imipridones in children with DMGs.

In summary, using a combination of metabolic imaging, mass spectrometry-based metabolomics, loss/gain-of-function studies, and clinically translatable therapeutic agents, we demonstrate that GABA production is a metabolic consequence of imipridone treatment that can be leveraged for therapy and for non-invasive metabolic imaging in preclinical DMG models.

## MATERIALS AND METHODS

### Cell culture

BT245 (male; RRID: CVCL_IP13) and SF8628 (female; RRID: CVCL_IT46) models were purchased from Sigma Aldrich and were originally isolated from pediatric patients harboring H3.3 K27M mutated tumors in the pons (*58–60*). Cells were transduced with firefly luciferase to enable optical imaging. The murine syngeneic 26-B7 and 26-C2 cell lines were a kind gift from Dr. Timothy Phoenix (*61, 62*). They were originally generated via *in utero* electroporation of PiggyBac DNA plasmids (expressing H3.3 K27M for 26-B7 and H3.1 K27M for 26-C3, dominant negative TP53 and mutant PDGFA) into the brainstem of pregnant C57BL/6 embryos at embryonic day 13.5. Once tumors developed, transplantable allograft cultures were established. 26-B7 and 26-C2 cells also express firefly luciferase and green fluorescent protein. All cell lines were maintained as neurospheres in Neurobasal-A medium (Gibco) containing 1% GlutaMAX, supplemented with 2% B27 (Gibco), 1% N2 (Gibco), 20 ng/mL human EGF, 20 ng/mL and antibiotics. Cell lines were routinely tested for mycoplasma contamination, authenticated by short tandem repeat fingerprinting (Cell Line Genetics) and assayed within 6 months of authentication.

### Viability, apoptosis and synergy assays

IC50 was measured by quantifying the percentage of live cells using a CellTiter-Fluor™ Cell Viability Assay (Promega) according to manufacturer’s instructions. Briefly, cells were seeded at 10,000 cells per well in a 96-well plate and treated with a 10-point serial dilution of ONC201 (Selleck Chemicals) or ONC206 (MedChemExpress) for 72 h. The CellTiter-Fluor™ reagent, which is cleaved to produce a fluorescent substrate only in live cells with protease activity, was then added and fluorescence measured at 400 nm/505 nm after 30 minutes at 37°C. IC50 was calculated by non-linear regression using GraphPad Prism 10. A similar assay format was used to measure apoptosis with the Caspase-Glo® 3/7 3D Assay (Promega). Cells were seeded and treated as described above for 72 h, following which the Caspase-Glo® 3/7 3D Reagent was added, and luminescence measured after 30 minutes at 37°C. To assess synergy, 10,000 cells/well were seeded in a 96-well plate and treated with a 5-point serial dilution of ONC206 or SGS-742 (MedChemExpress). Apoptotic cell count was measured as described above using the Caspase-Glo® 3/7 3D Assay (Promega). Bliss synergy scores were calculated using the SynergyFinder web application (version 3.0)61.

### Activity assays

GABA levels in cell pellets, culture medium and tumor tissue were measured using a human or mouse GABA ELISA kit (Biomatik) according to manufacturer’s instructions. Cell pellets were lysed by repeated freeze-thawing in saline. Tumor tissue was homogenized in PBS and cell culture supernatant was used after 1:1000 dilution in saline. MitoSOX Red, which is specific for superoxide radicals, was used to quantify superoxide radical generation in live cells as described earlier (*63*). Briefly, cells were seeded at 10,000 cells per well in a 96-well plate and treated with drugs or siRNA as needed for 72 h. Cells were washed and resuspended in saline and 5 μM MitoSOX Red was added to each well. After incubation at 37°C for 30 minutes, fluorescence was measured at 480 nm/570 nm. SOD1 activity was measured using a kit (Abcam, # ab65354) according to manufacturer’s instructions.

### Quantitative PCR (QPCR)

Gene expression was measured by QPCR and normalized to β-actin. The SYBR Green QPCR kit (Sigma) was used with the following primers: human *GAD1* (forward: GCGGACCCCAATACCACTAAC; reverse: CACAAGGCGACTCTTCTCTTC), mouse *GAD1* (forward: TCCAAGAACCTGCTTTCCTGT; reverse: GGATATGGCTCCCCCAGGAG), human *ABAT* (forward: GCTGAAATACCCTCTGGAAGAGT; reverse: AAGTCATCGGATGCGTGGTTG), mouse *ABAT* (forward: CAGGTGTTGAAGATCCGGTAG; reverse: CAGCAGACGTGATGACCTTC), human *SOD1* (forward: TCGTCTTGCTCTCTCTGGTC; reverse: CAGGCCTTCAGTCAGTCCTT), mouse *SOD1* (forward: AACCAGTTGTGTTGTCAGGAC; reverse: CCACCATGTTTCTTAGAGTGAGG), and β-actin (forward primer: AGAGCTACGAGCTGCCTGAC; reverse primer: AGCACTGTGTTGGCGTACAG).

### Silencing and overexpression studies

To silence ClpP, ATF4, GABBR1, or SOD1, lentiviral particles were generated by transfecting 293T cells with mouse or human ClpP sgRNA CRISPR/Cas9 All-in-One Lentivector plasmids (Abmgood) expressing a set of 3 sgRNAs against ClpP or ATF4 along with the third-generation packaging system using Lipofectamine 3000. DMG cells were then transduced with the lentiviral particles using polybrene and selected using puromycin. Cells transfected with a vector expressing scrambled sgRNA were used as controls. To overexpress ABAT or ATF4, cells were transiently transfected with the pCMV6-ENTRY-ABAT or pCMV6-ENTRY-ATF4 plasmid for 72 h using Lipofectamine 3000. Cells transfected with the pCMV6-ENTRY-control vector were used as controls. In all cases, gene expression was verified at 72 h by QPCR.

### ChIP-QPCR

Chromatin was immunoprecipitated using an antibody specific to ATF4 (Cell Signaling, #11815) or rabbit IgG (Cell Signaling, #2729) and the High-Sensitivity ChIP Kit (Abcam, ab185913) according to manufacturer’s instructions. QPCR was then performed for ABAT or GAD1 using the primers described above. Data was expressed as fold enrichment relative to IgG control.

### Western blots

Cells (∼10^7^) or ∼10 mg of tissue were lysed by sonication in RIPA buffer (25mM Tris-HCl pH 7.6, 150mM NaCl, 1% NP-40, 1% sodium deoxycholate, 0.1% SDS; Thermo Scientific) containing 0.5 mM phenylmethyl sulphonyl fluoride, 150 nM aprotinin and 1 μm each of leupeptin and E64 protease inhibitor. Lysates were cleared by centrifugation at 14000 rpm for 15 min at 4 °C and boiled in SDS-PAGE sample buffer (95°C for 10 min). Total cellular protein (∼20 µg) was separated on a 4-20% polyacrylamide gel (Bio-Rad) by sodium dodecyl sulphate polyacrylamide gel electrophoresis and transferred onto Immobilon-FL PVDF membrane (Millipore). Membranes were blocked overnight in 5% milk (Santa Cruz Biotechnology) in TBST (20 mM Tris-HCl, pH 7.5, 500 mM NaCl, 0.1% Tween 20) at 4°C. Membranes were then washed 3 times for 5 min each in TBST and incubated with primary antibodies diluted in TBST for 1 h at room temperature. Following 3 washes of 10 min each with TBST, HRP-conjugated secondary antibodies were added for 1 h in TBST at room temperature. Membranes were washed thrice in TBST for 10 min each and developed onto autoradiographic film using an enhanced chemiluminescence substrate kit (Thermo Scientific). Blots were probed for ClpX (Invitrogen, JE62-96), complex I (Cell Signaling, #70264), ATF4 (Cell Signaling, #11815), ABAT (Abcam, # ab108249), GAD1 (Cell Signaling, #41318), and SOD1 (Cell Signaling, #37385). β-actin (Cell Signaling, #4970) and α-actinin (Cell Signaling, #6487) were used as loading controls.

### MRS of cell extracts

Metabolites were extracted by the methanol-chloroform method (*64*). Briefly, cells were centrifuged, and washed with ice-cold saline. 10 ml each of ice-cold methanol (Sigma-Aldrich), chloroform (Acros Organics) and Milli-Q water were added with each step being followed by vortexing. Phase separation was achieved by centrifugation at 3000 rpm at 4 °C for 10 min, and solvents removed by lyophilization. The aqueous phase was reconstituted in 500 μl deuterium oxide (Acros Organics). ^1^H-MR spectra were obtained using *noesypr1d* sequence on a 600 MHz spectrometer (Bruker BioSpin) over 1h 4min. Peak integrals were quantified using MestReNova, corrected for saturation and normalized to an external reference (sodium 3-(trimethylsilyl) propionate-2,2,3,3-d4 (TSP; Sigma-Aldrich)) and to cell number (*64*).

### Metabolomics and stable isotope tracing in cells

∼5×10^5^ cells were seeded in T-25 flasks and treated with vehicle (DMSO), ONC206 (500 nM), or ONC201 (10 μM) in regular cell culture media for 72 h. For stable isotope tracing, cells were seeded and treated as above and incubated in media in which glutamine or glucose was replaced with [U-^13^C]-glutamine (99% enrichment; Cambridge Isotope Laboratories, final concentration 5.7 mM) or [U-^13^C]-glucose (99% enrichment; Cambridge Isotope Laboratories, final concentration 25 mM). Following incubation for 72 h, cells were washed with ice-cold ammonium acetate (150 mM, pH 7.3). 1 ml of pre-cooled methanol/water (80:20 v/v) was added to each well and plates incubated at -80 °C for 30 min. Cells were collected by scraping and centrifuged at 14,000 rpm for 15 min at 4 °C to remove debris. The supernatant was lyophilized, samples were reconstituted with 60 μL of pre-chilled acetonitrile/water (50:50, v/v), transferred into glass vials and utilized for liquid chromatography mass spectrometry (LC-MS) as described below.

### LC-MS

LC-MS was performed using a Vanquish Ultra High-performance LC system coupled to an Orbitrap ID-X Tribrid mass spectrometer (Thermo Fisher), equipped with a heated electrospray ionization (H-ESI) source capable of both positive and negative modes simultaneously (*65*). Before analysis, the MS instrument was calibrated using calibration solution (FlexMix, Thermo Fisher). Cell or tissue samples along with blank controls were placed in the autosampler. Individual samples were run alongside a pooled sample made from an equal mixture of all individual samples to ensure chromatographic consistency. Chromatographic separation of metabolites was achieved by hydrophilic interaction liquid chromatography (HILIC) using a Luna 3 NH2 column (150 mm x 2.1 mm, 3 μm, Phenomenex) in conjunction with a HILIC guard column (Phenomenex, 2.1 mm). The column temperature and flow rate were maintained at 27°C and 0.2 mL/min respectively. Mobile phases consisted of A (5 mM ammonium acetate, 48.5 mM ammonium hydroxide pH 9.9) and B (100% Acetonitrile). The following linear gradient was applied: 0.0-0.1 min: 85-80% B, 0.1-17.0 min: 80-5% B, 17.0-24.0 min: 5% B, 24.0-25.0 min: 5-85% B, 25.0-36.0 min: 85% B. The injection volume and auto sampler temperature were kept at 5 μL and 4°C respectively. High-resolution MS was acquired using a full scan method alternating between positive and negative polarities (spray voltages: +3800kV/-3100kV; sheath gas flow: 45 arbitrary units: auxiliary gas flow: 15 arbitrary units; sweep gas flow: 1 arbitrary unit; ion transfer tube temperature: 275°C; vaporizer temperature: 300°C). Three mass scan events were set for the duration of the 36-minute run time. The first was the negative polarity mass scan settings at full-scan-range; 70-975 m/z. Positive polarity mass scan settings were split to two scan events, full-scan-range; 70-360 m/z and 360-1500m/z, in that order. MS1 data were acquired at resolution of 60,000 with a standard automatic gain control and a maximum injection time of 100 ms. Data was acquired using Xcalibur software. Chromatograms were reviewed using FreeStyle (Thermo Fisher) and a 5-ppm mass tolerance. Peak areas were quantified using TraceFinder from either positive or negative modes depending on previously run standards. Peak areas were corrected to blank samples and normalized to cell number or wet weight of tissue and the total ion count of that sample. For ^13^C isotope tracing, % enrichment for each isotopomer was calculated after correcting for natural abundance using Escher Trace 63.

### Intracranial tumor implantation

All studies were performed in accordance with the National Institutes of Health Guide for the Care and Use of Laboratory Animals and were approved by the University of California, San Francisco Institutional Animal Care and Use Committee. 26B7, SF8628 or BT245 cells were intracranially injected into the cortex of C57BL/6 (for the 26B7) or SCID (for SF8628 and BT245) mice (female, 5-6 weeks old, Charles River Laboratories) as described previously (*64, 66*). BT245 cells were also injected into the pons of SCID mice using the following coordinates from the lambda suture (x=0.8 mm, y=6.5 mm, z=1 mm). For drug studies, mice were randomized and treated with vehicle (saline), ONC206 (25 mg/kg in saline, twice daily, 5 days per week) or the combination of ONC206 (25 mg/kg in saline, twice daily, 5 days per week) and SGS-742 (8 mg/kg, dissolved in 6.6% DMSO in saline, twice daily, 5 days per week) via intraperitoneal injection.

### *In vivo* MRI/^1^H-MRS studies

MR studies were performed using a ^1^H quadrature volume coil on a preclinical 3T MR scanner (Biospec, Bruker). Axial T2-weighetd images were acquired using a spin-echo TurboRARE sequence (TE/TR = 64/3700ms, FOV = 30 × 30mm^2^, 256 × 256, slice thickness = 1.5mm, NA = 5) (*66*). ^1^H-MRS spectra were acquired from a 27 mm^3^ voxel (for mice with cortical tumors) or 8 mm3 voxel (for mice with pontine tumors) placed in the center of the tumor using a point resolved spectroscopy (PRESS) sequence with the following parameters: TE = 16 ms (TE1 = TE2 = 8 ms), TR = 2000 ms, NA = 750, spectral width 2000 Hz and 2048 points (for the 27 mm^3^ voxel) or 1200 points (for the 8 mm^3^ voxel) combined with VAPOR water suppression. In addition, a spectrum of the unsuppressed water signal was acquired with the same parameters as above, but with a single average, for eddy current correction during LCModel spectral analysis. The spectral region between 0.2 and 4.0 ppm was analyzed using LCModel and a basis set comprised of GABA, creatine, glutathione, phosphocreatine, N-acetyl aspartate (NAA), N-acetyl aspartyl glutamate (NAAG), choline, lactate, glucose, phosphocholine, glyceryl-phosphocholine, phosphoethanolamine (PE), ascorbate, inositol, aspartate, taurine, alanine, glutamine, glutamate, glycine, scyllo-inositol. Each peak was normalized to the total signal calculated as the sum of all the quantified metabolites. The average concentration for each metabolite was calculated as before (*64*). Average metabolic changes were then calculated for each metabolite of interest.

### Optical imaging

Tumor growth was monitored by bioluminescence on a Xenogen IVIS Spectrum 10 min after intraperitoneal injection of 150 mg/kg D-luciferin prepared according to the manufacturer’s directions (Gold Biotechnology, #LUCK-100). The total flux signal was used for quantification in all studies.

### Statistical analysis

All experiments were performed on a minimum of 3 biological replicates (n≥3) and results presented as mean ± standard deviation. Statistical significance was assessed in GraphPad Prism 10 using a two-way ANOVA or two-tailed Welch’s t-test with p<0.05 considered significant. Analyses were corrected for multiple comparisons using Tukey’s method, wherever applicable. * indicates p<0.05, ** indicates p<0.01, *** indicates p<0.001 and **** indicates p<0.0001.

## Author contributions

Conceptualization: PV, SM, SV, CK

Methodology: PV, SV, CK, GB, SU, AMG, CT, BL, SJ, TP

Investigation: PV, SV, CK, GB, SU, AMG, CT, BL, SJ, TP

Visualization: PV, GB

Funding acquisition: PV, SM, GB

Project administration: PV, CK

Supervision: PV, CK

Writing – original draft: PV, GB

Writing – review & editing:

## Data and materials availability

This manuscript does not report any original code. All data generated in this study are available in the main text or the supplementary materials. Any additional information required to reanalyze the data reported in this manuscript is available from the corresponding author upon reasonable request.

