## Supplementary Figures 1-6 for "GABA production induced by imipridones is a targetable and imageable metabolic alteration in diffuse midline gliomas"

SUPPLEMENTARY FIGURE 1

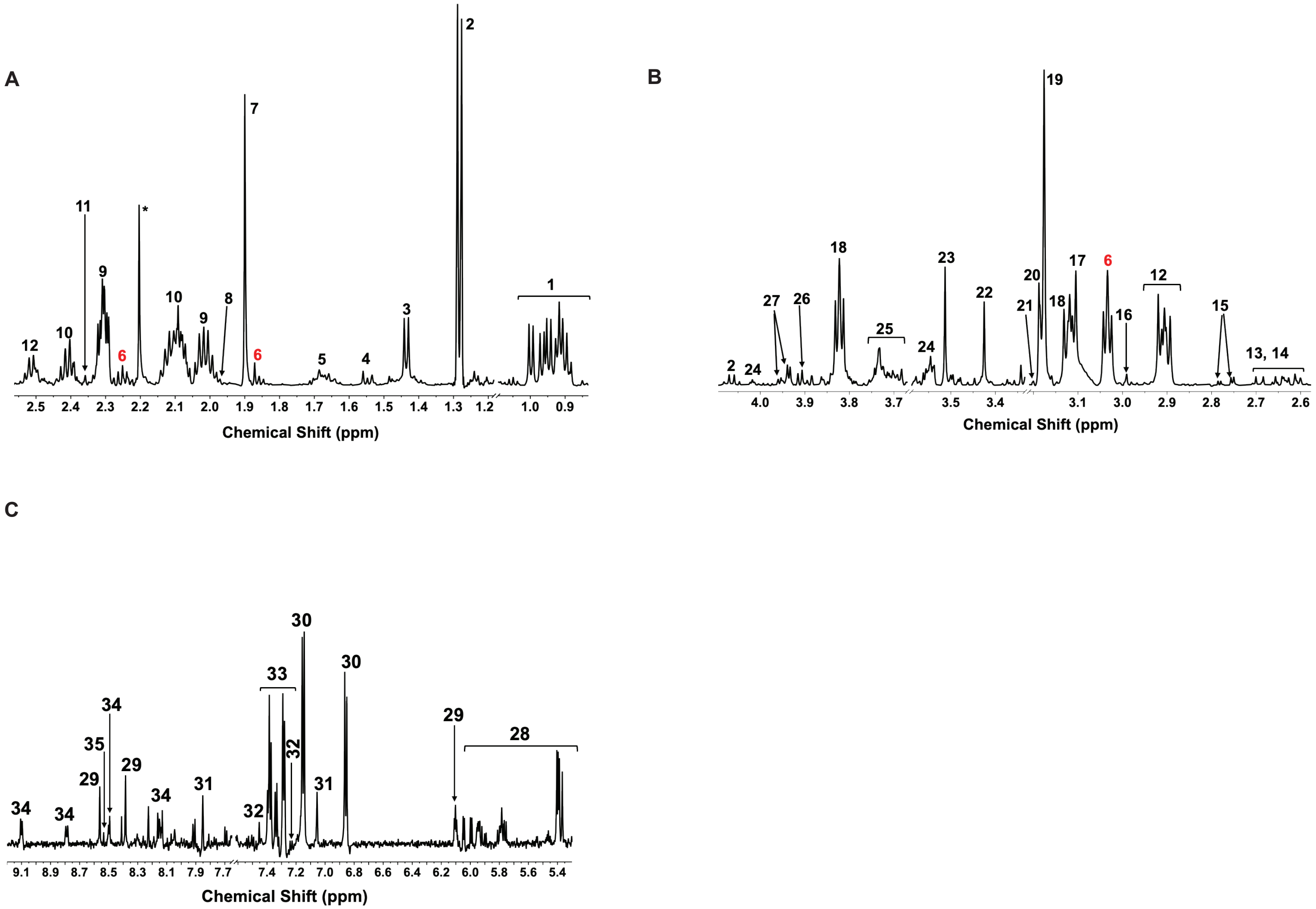

### Supplementary Figures

**Supplementary Fig. 1**. **$^{1}$H-MRS identifies GABA as a unique metabolic biomarker in patient-derived and syngeneic DMG cells.** Representative $^{1}$H-MRS spectra from BT245 cells treated with 500 nM ONC206 for 72 h. Panel A shows peak assignments for the spectral region from 0.8 to 2.6 ppm, panel B shows the spectral region from 2.6 to 4.2 ppm and panel C shows the spectral region from 5.3 to 9.2 ppm. Assignments: 1: Branched chain amino acids; 2: lactate/threonine; 3: alanine; 4: lipids; 5: lysine, arginine, ornithine; 6: GABA; 7: Acetate; 8: N-acetyl aspartate; 9: glutamate; 10: glutamine; 11: succinate; 12: glutathione; 13: Citrate; 14: Methionine; 15: Aspartate; 16: α-ketoglutarate; 17: Creatine; 18: Phosphocreatine; 19: choline; 20: phosphocholine; 21: glycerophosphocholine; 22: taurine; 23: glycine; 24: myoinositol; 25: serine, glucose, UDP-glucose, UDP-N-acetylglucosamine, UDP-glucuronate. 26: serine; 27: fructose; 28: galactose-1-phosphate, UDP-N-acetylglucosamine, UDP-glucose, UDP-glucuronate, UMP, NAD; 29: AXP; 30: tyrosine; 31: histidine; 32: tryptophan; 33: phenylalanine; 34: NAD+; 36: formate.

SUPPLEMENTARY FIGURE 2

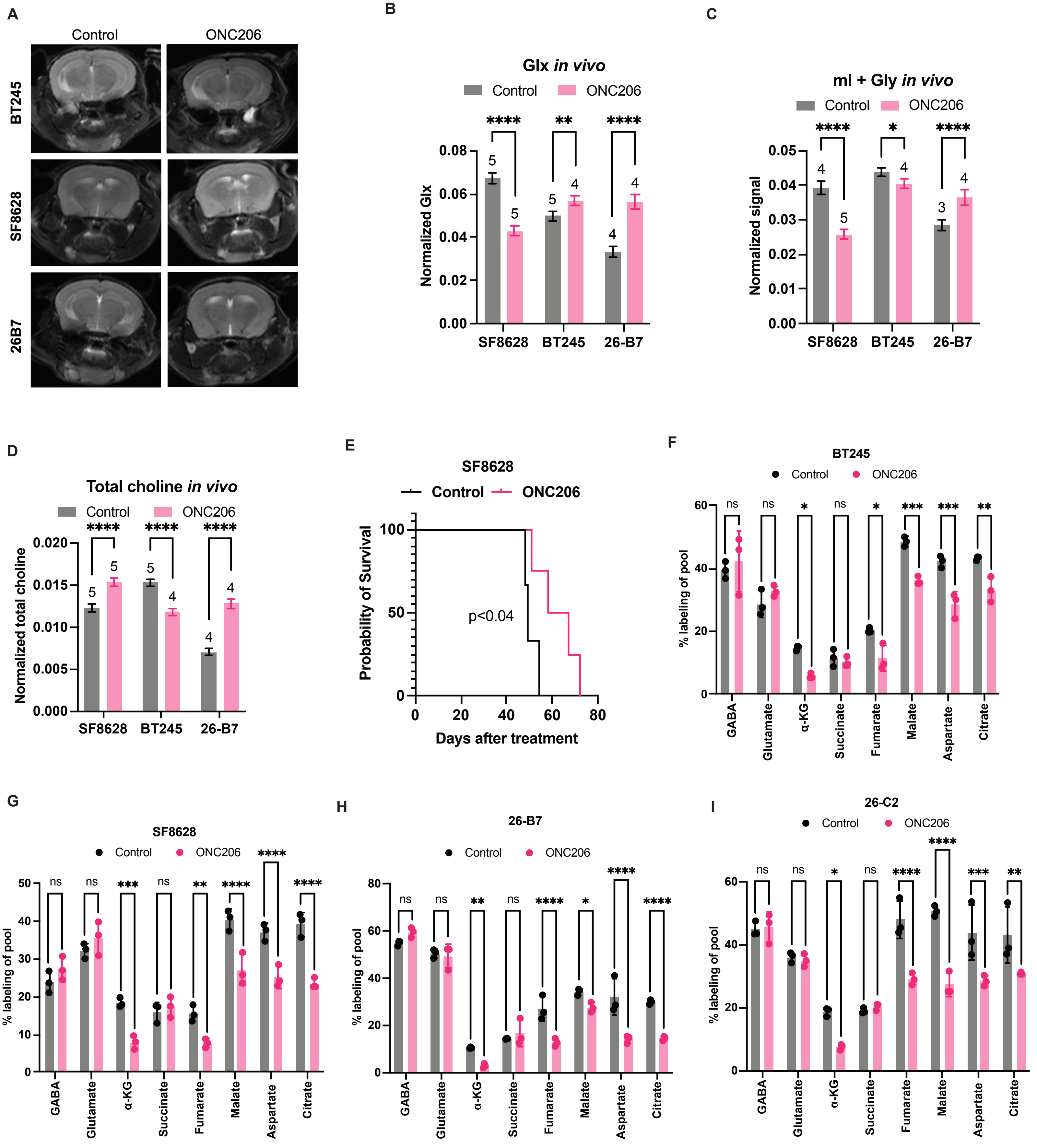

**Supplementary Fig. 2**. **$^{1}$H-MRS-detectable GABA is an early imaging biomarker of drug-target engagement in mice bearing orthotopic DMG xenografts *in vivo.*** **(A)** Representative T2-weighted MRI from mice bearing intracranial BT245, SF8628, or 26-B6 xenografts treated with vehicle (saline) or ONC206 (25 mg/kg, twice daily). Data was acquired on day 7±1 on a Bruker 3T scanner. Quantification of *in vivo* $^{1}$H-MRS-detectable glutamate + glutamine (glx) **(B)**, myoinositol + glycine (mI + Gly, **C**), or total choline (sum of choline, phosphocholine, glycerophosphocholine, **D**) from mice bearing orthotopic BT245, SF8628, or 26-B7 tumors treated with vehicle (saline) or ONC206 (25 mg/kg, twice daily). On day 7±1, $^{1}$H-MRS data was acquired on a Bruker 3T scanner. **(E)** Survival curves for mice bearing intracranial SF8628 xenografts treated with vehicle (saline) or ONC206 (25 mg/kg, twice daily, 5 days/week). Mice were monitored until they had to be euthanized according to IACUC guidelines. Quantification of % $^{13}$C labeling of metabolites from [U-$^{13}$C]-glucose in BT245 **(F)**, SF8628 **(G)**, 26-B7 **(H)** and 26-C2 **(I)** cells treated with vehicle (DMSO) or 500 nM ONC206 for 72 h.

SUPPLEMENTARY FIGURE 3

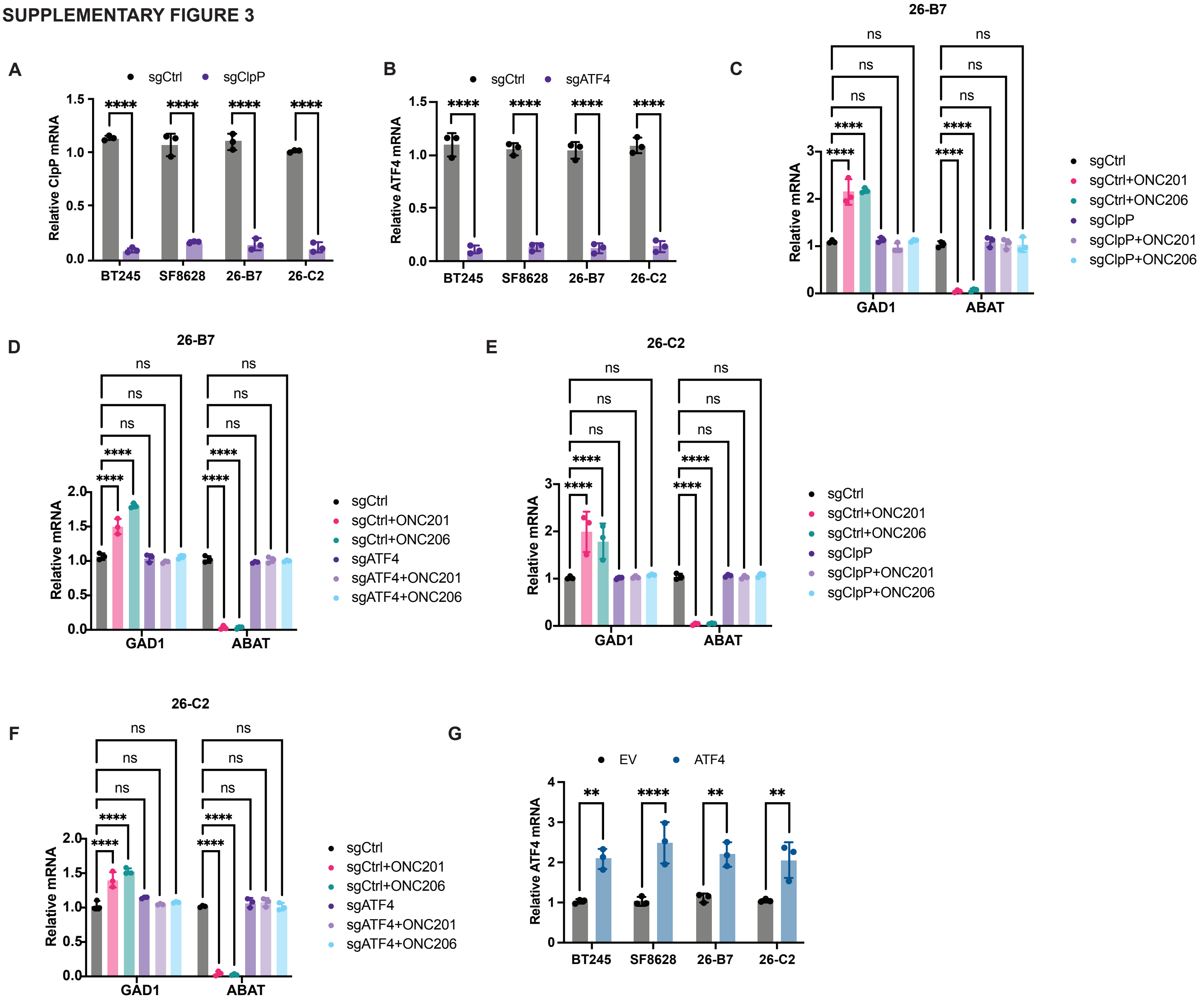

**Supplementary Fig. 3**. **Imipridones upregulate GAD1 and downregulate ABAT via ClpP and ATF4 in DMG cells.** Verification of ClpP **(A)** and ATF4 **(B)** silencing by pooled sgRNA in BT245, SF8628, 26-B7 and 26-C2 cells. ClpP and ATF4 mRNA were measured by QPCR. Expression of GAD1 and ABAT measured by QPCR in 26-B7 cells expressing sgRNA against ClpP **(C)** or ATF4 **(D)**. Cells expressing a scrambled sgRNA sequence were used as control (sgControl). Cells were treated with vehicle (DMSO), 500 nM ONC206, or 10 μM ONC201 for 72 h. Expression of GAD1 and ABAT measured by QPCR in 26-C2 cells expressing sgRNA against ClpP **(E)** or ATF4 **(F)**. Cells expressing a scrambled sgRNA sequence were used as control (sgControl). Cells were treated with vehicle (DMSO), 500 nM ONC206, or 10 μM ONC201 for 72 h. **(G)** Confirmation of increased ATF4 expression in BT245, SF8628, 26-B7 and 26-C2 cells expressing an empty vector (EV) or a plasmid expressing ATF4 (ATF4).

SUPPLEMENTARY FIGURE 4

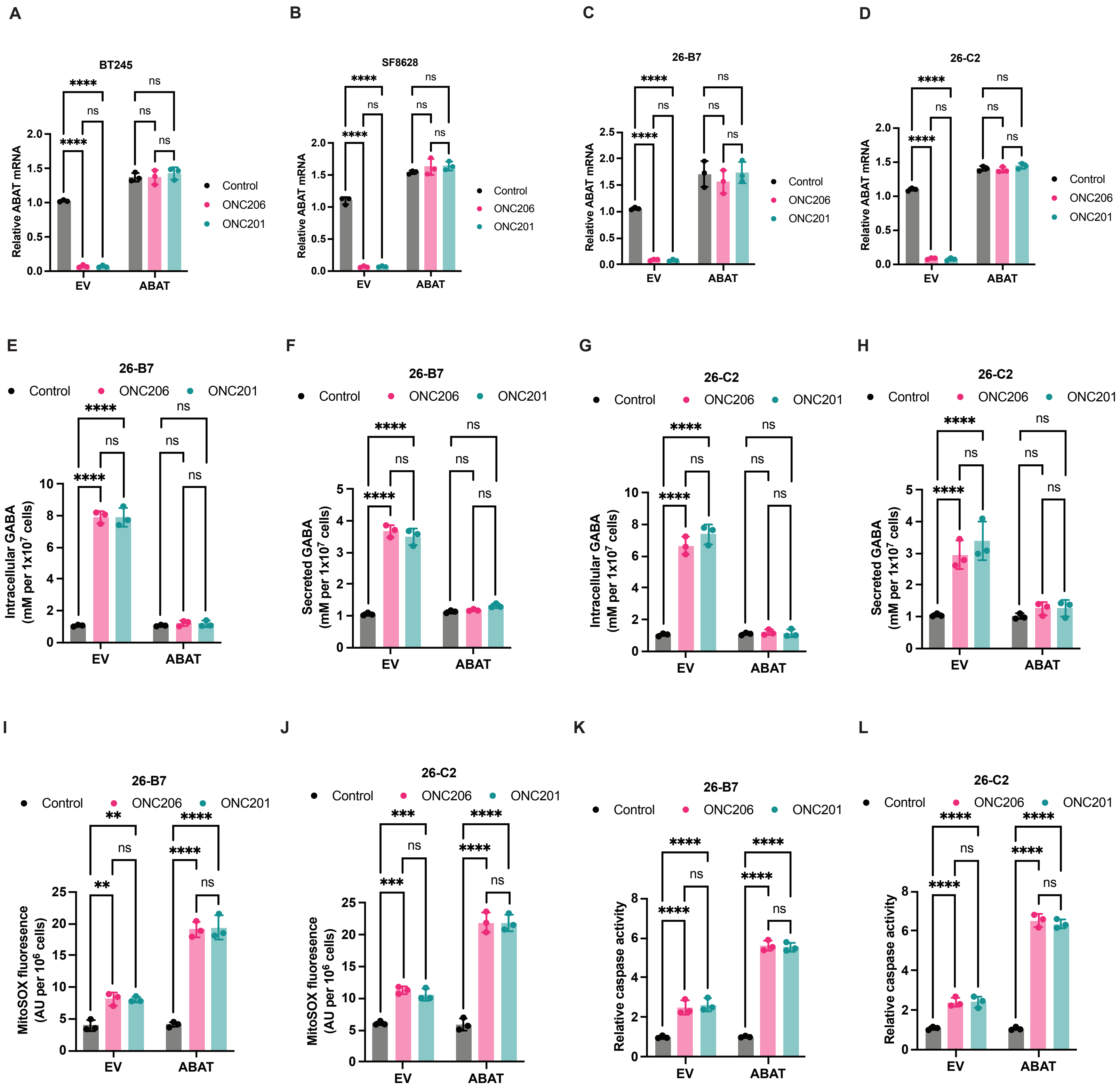

**Supplementary Fig. 4**. **Depleting GABA by ABAT overexpression potentiates superoxide radical generation and apoptosis in imipridone-treated DMG cells.** Verification of ABAT overexpression in BT245 **(A)**, SF8628 **(B)**, 26-B7 **(C)**, and 26-C2 **(D)** cells expressing an empty vector (EV) or a plasmid expressing ABAT (ABAT). Cells were treated with vehicle (DMSO), 500 nM ONC206, or 10 μM ONC201 for 72 h. Intracellular **(E)** and secreted **(F)** GABA concentration in 26-B7 cells transfected with an empty vector (EV) or a plasmid expressing ABAT (ABAT). Cells were treated with vehicle (DMSO), 500 nM ONC206, or 10 μM ONC201 for 72 h. Intracellular **(G)** and secreted **(H)** GABA concentration in 26-C2 cells transfected with an empty vector (EV) or a plasmid expressing ABAT (ABAT). Cells were treated with vehicle (DMSO), 500 nM ONC206, or 10 μM ONC201 for 72 h. Superoxide radical levels in 26-B7 **(I)** and 26-C2 **(J)** cells transfected with an empty vector (EV) or a plasmid expressing ABAT (ABAT). Cells were treated with vehicle (DMSO), 500 nM ONC206, or 10 μM ONC201 for 72 h. Caspase activity measured using the Caspase-Glo® 3/7 3D Assay in 26-B7 **(K)** and 26-C2 **(L)** cells transfected with an empty vector (EV) or a plasmid expressing ABAT (ABAT). Cells were treated with vehicle (DMSO), 500 nM ONC206, or 10 μM ONC201 for 72 h.

SUPPLEMENTARY FIGURE 5

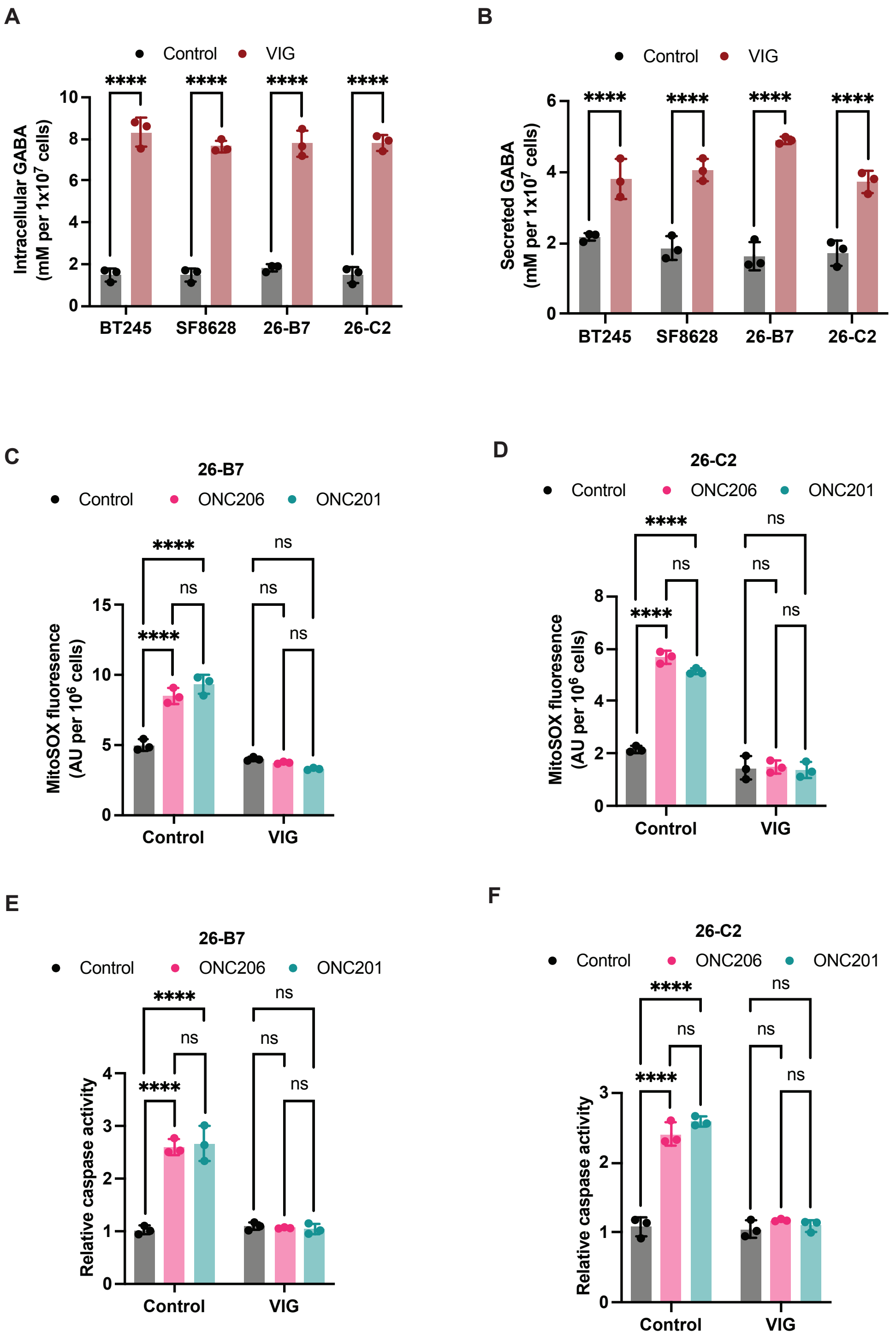

**Supplementary Fig. 5**. **Enhancing GABA accumulation by treatment with the ABAT inhibitor vigabatrin potentiates superoxide radical generation and apoptosis in imipridone-treated DMG cells.** Intracellular **(A)** and secreted **(B)** GABA concentration in BT245, SF8628, 26-B7 and 26-C2 cells treated with vehicle (DMSO) or vigabatrin (100 μM). Cells were concurrently treated with vehicle (DMSO), 500 nM ONC206, or 10 μM ONC201 for 72 h. Superoxide radical levels in 26-B7 **(C)** and 26-C2 **(D)** cells treated with vehicle (DMSO) or vigabatrin (100 μM). Cells were concurrently treated with vehicle (DMSO), 500 nM ONC206, or 10 μM ONC201 for 72 h. Caspase activity measured using the Caspase-Glo® 3/7 3D Assay in 26-B7 **(E)** and 26-C2 **(F)** cells treated with vehicle (DMSO) or vigabatrin (100 μM). Cells were concurrently treated with vehicle (DMSO), 500 nM ONC206, or 10 μM ONC201 for 72 h.

### SUPPLEMENTARY FIGURE 6

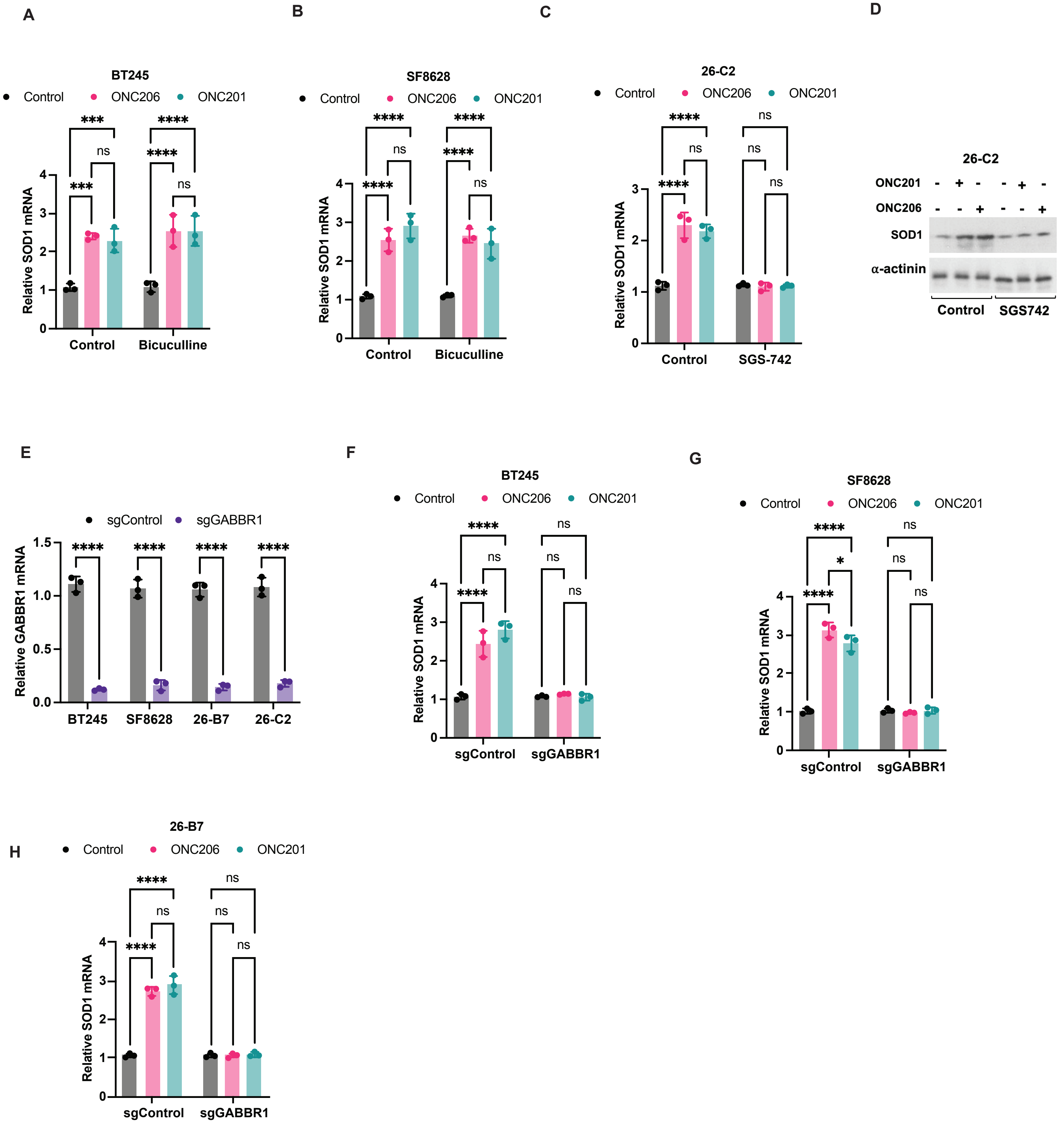

**Supplementary Fig. 6**. **Autocrine GABA signaling via the GABAB receptor drives SOD1 expression in imipridone-treated DMG cells.** SOD1 mRNA in BT245 **(A)** and SF8628 **(B)** cells treated with vehicle (DMSO) or the GABAA receptor antagonist bicuculline (10 μM). Cells were concurrently treated with vehicle (DMSO), 500 nM ONC206, or 10 μM ONC201 for 72 h. SOD1 mRNA **(C)** and protein **(D)** in 26-C2 cells treated with vehicle (DMSO) or the GABAB receptor antagonist SGS-742 (50 μM). Cells were concurrently treated with vehicle (DMSO), 500 nM ONC206, or 10 μM ONC201 for 72 h. **(E)** Verification of GABBR1 silencing in BT245, SF8628, 26-B7 and 26-C2 cells expressing scrambled sgRNA (sgControl) or GABBR1-targeted pooled sgRNA (sgGABBR1). GABBR1 mRNA was measured by QPCR. SOD1 mRNA in BT245 **(F)**, SF8628 **(G)**, and 26-B7 **(H)** cells expressing scrambled sgRNA (sgControl) or GABBR1-targeted pooled sgRNA (sgGABBR1). SOD1 mRNA was measured by QPCR.
